# Dilp8 relaxin signaling from ovarian follicle cells to Lgr3+ neurons promotes spontaneous ovulation and oocyte quality in *Drosophila*

**DOI:** 10.1101/2025.04.03.647069

**Authors:** Yanel Volonté, Fabiana Heredia, Rebeca Zanini, Juliane Menezes, María Sol Perez, Mafalda Gualdino, Lara Lage, Leonor da Silva Luz, Andreia P. Casimiro, Catarina. C.F. Homem, Maria Luísa Vasconcelos, Andres Garelli, Alisson M. Gontijo

**Author notes:** These authors contributed equally to this work (Yanel Volonté, Fabiana Heredia, Rebeca Zanini).

## Abstract

Ovulation enables mature oocytes to exit the ovary for potential fertilization. In *Drosophila*, ovulation is induced by mating but also occurs spontaneously in virgins, with rates varying widely in natural populations: short oocyte retention is ancestral, while longer retention is favored in colder climates. The molecular regulation of spontaneous ovulation remains unclear. Here, we show that disrupting the relaxin/insulin-like peptide Dilp8 or its receptor Lgr3—an orthologue of vertebrate RXFP1/2—in follicle cells or specific neurons, respectively, delays ovulation, slows average egg transit time in the reproductive tract, and facilitates oogenesis progression beyond ∼2 mature oocytes per ovariole, leading to mature follicle accumulation in the ovary. Mating largely rescues these defects, suggesting the pathway is dispensable post-mating. Dilp8-Lgr3 signaling ensures high oocyte quality by promoting elimination of lower-quality aging oocytes and by antagonizing oogenesis progression via an undefined mechanism downstream of Lgr3+ neurons. Our findings provide a molecular basis for oocyte retention time regulation in *Drosophila* involving ovarian-nervous system cross-talk, and bring further support for an ancient, conserved role for relaxin-like signaling in regulating ovulation and overall female reproductive physiology.

**Summary statement:** This study uncovers how ovary–neuron communication promotes spontaneous ovulation and prevents buildup of aging eggs, ensuring optimal egg quality in fruit flies.

## INTRODUCTION

Ovulation is a fundamental biological process in which a mature oocyte sheds its surrounding follicle cells and leaves the ovary, enabling the possibility of fertilization (Bloch Qazi et al., 2003; Richards 2007). Ovulation can be influenced by environmental and internal states, such as seasons, lunar cycles, stress, pheromones, mating status, and pregnancy in ways that are many times species-specific (Bloch Qazi et al., 2003; Brown, 2018;; Richards, 2007; Mercier and Hamel 2009; Rekwot et al., 2001; Yamamoto et al., 2006). In *Drosophila*, like in other species, ovulation is largely triggered by mating. This occurs via at least two processes: (1) a seminal protein, ovulin (Bloch Qazi et al., 2003; Heifetz et al., 2000), increases ovulation rates in a process that is dependent on octopamine (Rubinstein and Wolfner, 2013), and (2) female spermathecal secretory cell factors also contribute to increasing postmating ovulation rate (Schnakenberg et al., 2011; Sun and Spradling, 2013).

During sexual maturation, virgin *Drosophila* females accumulate mature oocytes and ovulate significantly less than mated females throughout their lives (Boulétreau-Merle, 1990; Boulétreau-Merle and Fouillet, 2002). The timing of the onset of virgin ovulation, also called oocyte retention, is a life-history trait that varies clinally and seasonally in natural populations and has a genetic basis (Akhund-Zade et al., 2017; Boulétreau-Merle et al., 1992; Horváth and Kalink,a 2018). Short retention phenotypes, where laying of unfertilized eggs starts as soon as 2 days after adult emergence, are considered the ancestral state of *Drosophila melanogaster*, while long retention phenotypes, where the first unfertilized eggs are laid up to 15 days after emergence, are considered adaptations to cold environments and overwintering (Boulétreau-Merle, 1990; Boulétreau-Merle and Fouillet 2002). Whereas many species, such as murines and primates, also ovulate spontaneously (Bakker and Baum 2000), energy investment in oogenesis is considered relatively higher for *Drosophila* females, so virgin ovulation is regarded as a residual, unregulated process and a pointless loss of reproductive potential (Akhund-Zade et al., 2017; Boulétreau-Merle and Fouillet, 2002; Horváth and Kalinka, 2018).

Drosophila insulin-like peptide 8 (Dilp8) is a relaxin-like peptide that promotes developmental stability and pupariation behavior progression during late larval stages (Colombani et al., 2012; Garelli et al., 2012; Gontijo and Garelli, 2018; Heredia et al., 2021). In both cases, Dilp8 acts via a conserved neuronal receptor, the Leucine rich repeat containing G protein coupled receptor 3, Lgr3, the orthologue of vertebrate RXFP1/2 relaxin receptors (Colombani et al., 2015; Garelli et al., 2015; Jaszczak et al., 2016; Vallejo et al., 2015). Dilp8 is highly expressed in the adult ovary (Chintapalli et al., 2007; Garelli et al., 2012), specifically in terminal-stage follicle cells coating mature oocytes (Li et al., 2022; Li et al., 2023; Liao and Nässel, 2020). Follicle-cell derived Dilp8 has been proposed to generally regulate fecundity, ovulation, and metabolism in adults (Li et al., 2023; Liao and Nässel, 2020). Endogenous *Lgr3* has also been proposed to regulate fecundity (Li et al., 2023). Moreover, a subset of sexually-dimorphic *Lgr3*-positive neurons has been shown to regulate female receptivity and fecundity (Meissner et al., 2016). Finally, expression of Lgr3 GAL4 lines was found in posterior follicle cells and in neurons innervating the oviduct, which could suggest that Dilp8 acts via both autocrine and paracrine mechanisms (Liao and Nässel, 2020). It is unclear, however, if the above-mentioned neuronal activities are mediated by or require *Lgr3* at all, and if the neurons that control fecundity are the same as those innervating the proximal oviduct.

Here, we show that follicle-cell-derived Dilp8 acts either paracrinally or systemically on at least two subpopulations of Lgr3+ neurons to promote ovulation and egg-laying control, but mostly in virgin females. Mating largely rescues these defects, suggesting the Dilp8-Lgr3 pathway is dispensable post-mating. In parallel, Dilp8 and Lgr3 also inhibit oogenesis progression, so that in their absence, virgins bypass a checkpoint that otherwise limits the number of mature follicles per ovariole to ∼2, leading to an accumulation of mature follicles in the ovary. Dilp8-Lgr3 pathway activity thus contributes to promoting optimal egg quality by eliminating aged oocytes by spontaneous ovulation and by quantitatively inhibiting oogenesis progression proportionally to the amount of mature, Dilp8+ follicles accumulated in the ovaries.

## RESULTS

### Dilp8 is expressed in and secreted from mature, stage 14, follicle cells

Previous work has shown that *dilp8* mRNA is highly enriched in the ovary of *D. melanogaster* and other flies, specifically in the follicle cells of stage 14 follicles (Garelli et al., 2012; Li et al. 2022; Li et al., 2023; Liao and Nässel, 2020; Tootle et al., 2011;). We confirmed these findings using the transcriptional reporter *dilp8-enhanced Green Fluorescent Protein (eGFP)^MI00727^*, which drives similar expression as the late-follicle-cell-specific driver *R47A04-GAL4* (Deady et al., 2015; Pfeiffer et al., 2008) (**Fig. 1A**), and the enhancer trap *dilp8-GAL4* line (*dilp8-1-GAL4*) (**Fig. 1A**), which are also loss of function alleles (**Fig. 1C**), and with an anti-Dilp8 antibody (Colombani et al., 2012) (**Fig. 1B**). The vesicular staining pattern found, comparable to that reported with an independent antibody (Liao and Nässel, 2020), suggests that Dilp8 could be continuously released from follicle cells, rather than upon follicle cell rupture during ovulation.

**Fig. 1.**
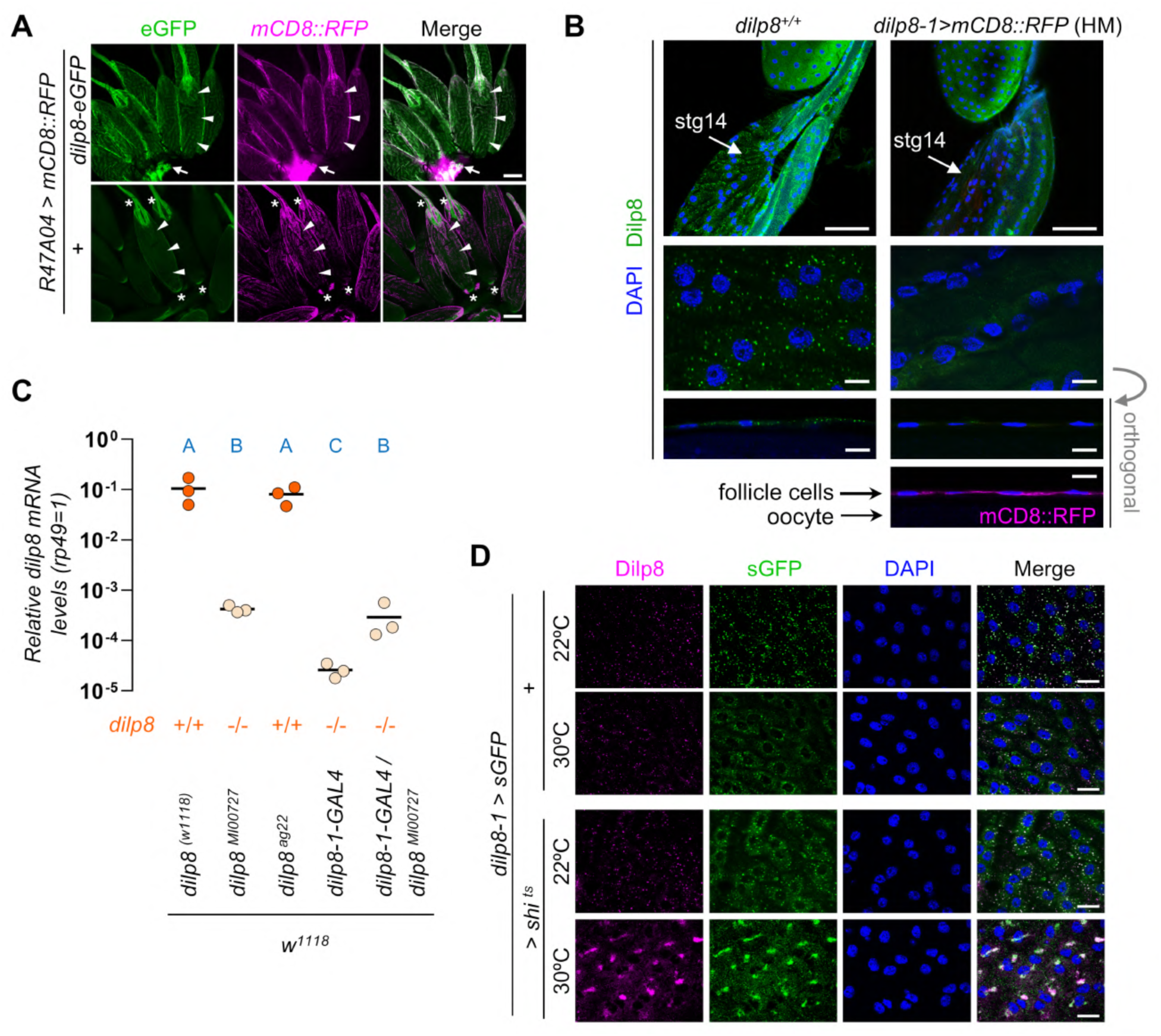
Dilp8 is expressed in and secreted from mature, stage 14 follicle cells. **(A)** Top panel: the transcriptional reporter *dilp8-eGFP[MI00727]* (green) is expressed in mature oocyte follicle cells (arrowheads) identified with the late-follicle-cell-specific driver *R47A04-GAL4* driving *UAS-mCD8::RFP* (magenta). The arrow points to the residual body. Bottom panel: animals not carrying *dilp8-eGFP[MI00727]* have no detectable eGFP expression. Asterisks mark autofluorescent signal. **(B)** Top panel, Dilp8 immunostaining shows a punctate pattern in stage 14 follicle cells that is absent in mutants carrying two copies of *dilp8-GAL4-1* [*dilp8-1>mCD8::RFP* (HM, for homozygous)]. Earlier follicles show unspecific fluorescence, also present in *dilp8* mutants. Middle and bottom panels, higher resolution images of stage 14 follicle cells. Bottom panels, orthogonal views of follicle cells. The top, middle, and bottom panels are not necessarily from the same follicles. **(C)** Expression levels of *dilp8* measured by quantitative reverse-transcriptase polymerase chain reaction (qRT-PCR) of the indicated genotypes. Dots: independent ovary samples. Same blue letter, *P* > 0.01, ANOVA and Tukey’s HSD test. **(D)** Flies that carry *shibire^ts^* (*>shi^ts^*) expressed in mature follicle cells with a single copy of *dilp8-GAL4-1* (*dilp8-1>*) lose the Dilp8 vesicular pattern upon shifting from 22 °C to 30 °C for one day. A constitutively secreted GFP (*sGFP*) protein was enriched in the same compartments as Dilp8 at both temperatures. Scale bar = 100 µm (A), 50 µm (B, top panels), 10 µm (B, middle and bottom panels, and C).

To test this hypothesis, we inhibited secretory vesicle recycling in stage 14 follicle cells of 7-day old flies by expressing the temperature-sensitive mutant of the Dynamin*-*encoding gene, *shibire^ts^* (*shi^ts^*) (Kitamoto, 2001) under the control of *dilp8-1-GAL4* (*dilp8>shi^ts^*) together with a constitutively secreted GFP (sGFP) (Entchev et al., 2000). Dynamin is required for secretory vesicle recycling and vesicle budding from the trans-Golgi network (Jones et al., 1998). The mutant dynamin allele *shi^ts^* blocks both processes at restrictive temperature and is expected to cause secretory vesicle depletion only if vesicles are effectively secreted during the temperature shift period. At the permissive temperature of 22°C, both Dilp8 and sGFP were found in cytoplasmic vesicles (**Fig. 1D**). After 24 h at the restrictive temperature (>29°C), the pool of constitutively secreted sGFP was depleted and sGFP was observable only in the perinuclear compartment.

The same pattern was found for Dilp8 (**Fig. 1D**). These findings are coherent with Dilp8 being secreted from the follicle cells in the ovary, while they are still surrounding mature oocytes. In this experiment, *dilp8>shi^ts^*flies or flies expressing *shi^ts^* in follicle cells using the *R47A04-GAL4* line (Deady et al., 2015; Pfeiffer et al., 2008) quantitatively lay drastically fewer unfertilized eggs than controls (**Fig. S1**), consistent with a model where constitutive Dilp8 secretion from stage 14 follicles is required for proper virgin ovulation.

### Dilp8 mutant virgins retain oocytes in the ovary

Virgins that are homozygous for the hypomorphic allele *dilp8-eGFP^MI00727^*and backcrossed to the *w^1118^* background (hereafter, *dilp8-eGFP^MI00727^*) have bloated abdomens relative to control *w^1118^* virgins (**Fig. 2A**) due to a large amount of late-stage oocytes in the ovary (**Figs 2A, S2)** and a lower number of unfertilized eggs laid (**Fig. 2B**). The precise excision of the *eGFP^MI00727^* cassette restored *dilp8* mRNA levels (**Fig. 1C**), and resulted in an increase in the number of unfertilized eggs laid as virgins (**Fig. 2B**), suggesting that the virgin egg retention phenotype is linked to Dilp8 levels. Strikingly, the reduced egg-laying phenotype was absent in mated females when comparing *dilp8-eGFP^MI00727^* with the *dilp8^ag22^* precise excision and partially rescued relative to *w^1118^* controls (**Fig. 2B**). This contrasts with previous reports suggesting that mated (and non-backcrossed) *dilp8-eGFP^MI00727^* mutant animals had reduced fecundity (Li et al., 2023; Liao and Nässel, 2020), suggesting that this discrepancy could be an effect of heterogeneous backgrounds. We found similar results as above with an independent *dilp8* null allele, *dilp8^ag50^*(Heredia et al., 2021), when compared to its wild-type background allele, *dilp8^ag52^*(**Fig. 2C**), or with RNA interference (RNAi)-mediated knockdown of *dilp8* [*UAS-dilp8-IR* (Dietzl et al., 2007; Perkins et al., 2015)] specifically in follicle cells (*R47A04>dilp8-IR*) (**Fig. 2D**). Again, even though a small significant effect on fecundity was detectable in mated *dilp8^ag50^*null alleles, the effect size (21% reduction) was visibly smaller than that of virgins (81% reduction). We conclude that follicle cell-derived Dilp8 positively influences the number of unfertilized eggs laid in virgin females, and that mating renders this influence largely redundant.

**Fig. 2.**
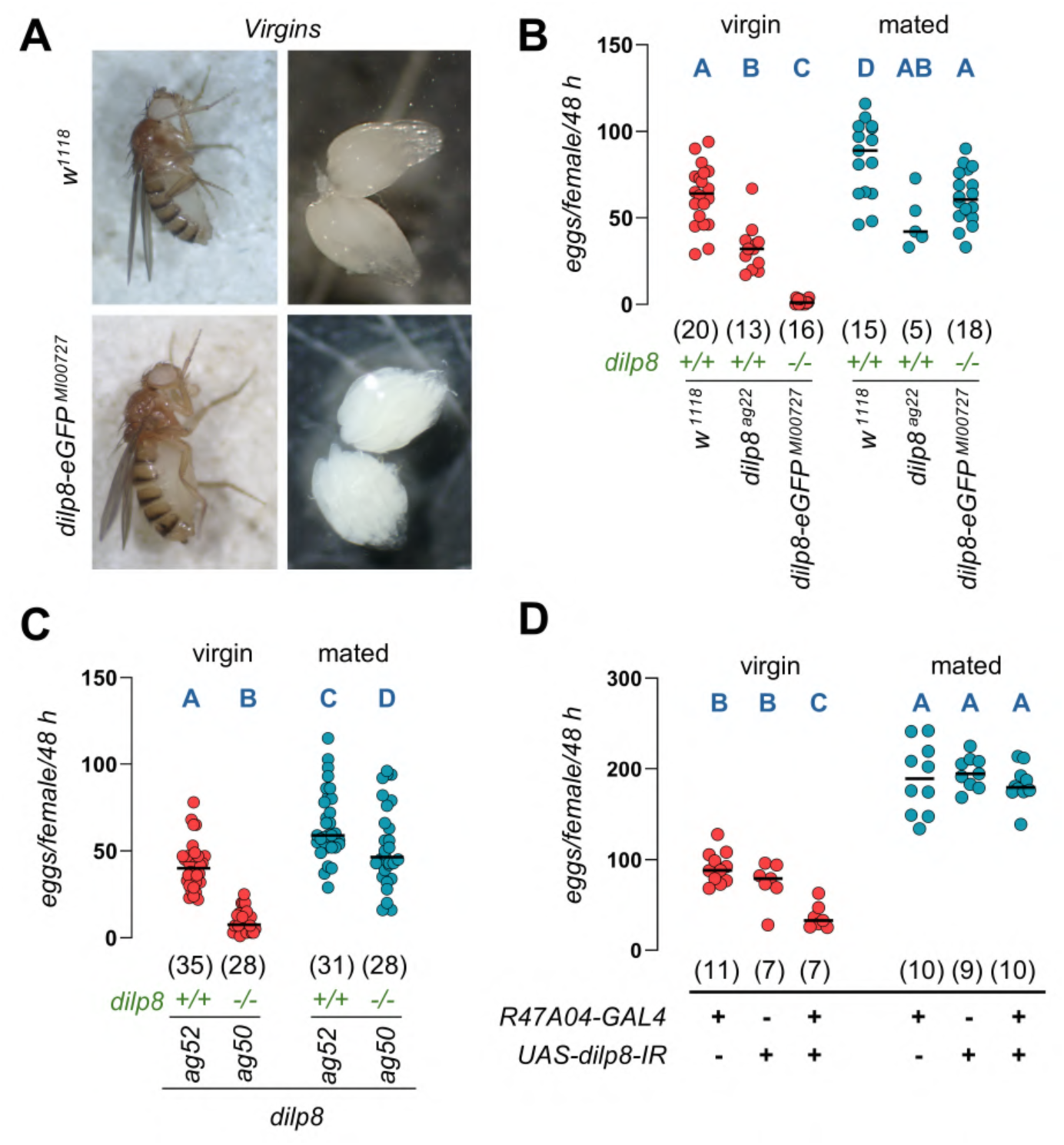
Dilp8 is required in virgin females for ovulation. **(A)** The abdomen of *dilp8* mutant flies was bloated due to the accumulation of eggs in the ovaries. **(B)** Virgin females lacking *dilp8* (*dilp8-eGFP^MI00727^)* laid fewer eggs than *w^1118^* and *dilp8^ag22^* control flies. The effect was lost upon mating. **(C)** Virgin females carrying a null allele of *dilp8* (*dilp8^ag50^*) laid fewer eggs than their background controls (*dilp8^ag52^*). The effect size is reduced upon mating. **(D)** Knockdown of *dilp8* in follicle cells reduces the number of eggs laid by virgin females but not mated females. *N* of biological replicates, each with five individuals, in parentheses. Same letter *P* > 0.05, ANOVA and Tukey’s HSD test (B, C) and Brown-Forsythe and Welch ANOVA and Dunnett’s T3 test (D).

The same results were observed when the number of eggs laid was evaluated for longer time periods. Unfertilized egg laying was constantly lower (1.4-2.7-fold) in *dilp8* mutants than in control *dilp8^ag52^*females evaluated for 10 days starting 5 days after eclosion (**Fig. 3A**). This difference was statistically significant in 9/10 days in virgins (**Fig. 3A**), and 0/10 days in mated females (**Fig. 3B**). Direct comparison of the fold reduction in egg laying caused by *dilp8* mutation in virgins and mated animals corroborated this specific reduction in virgin females (**Fig. 3C**), and revealed that the effect was highest in the three initial days (days 5-7 post eclosion), plateauing at ∼1.5 fold lower egg laying for the next 7 days (**Fig. 3C**). As *dilp8* mutants are capable of the same amount of egg laying as controls in mated conditions, these results indicate that Dilp8 plays no major role in promoting oogenesis.

**Fig. 3.**
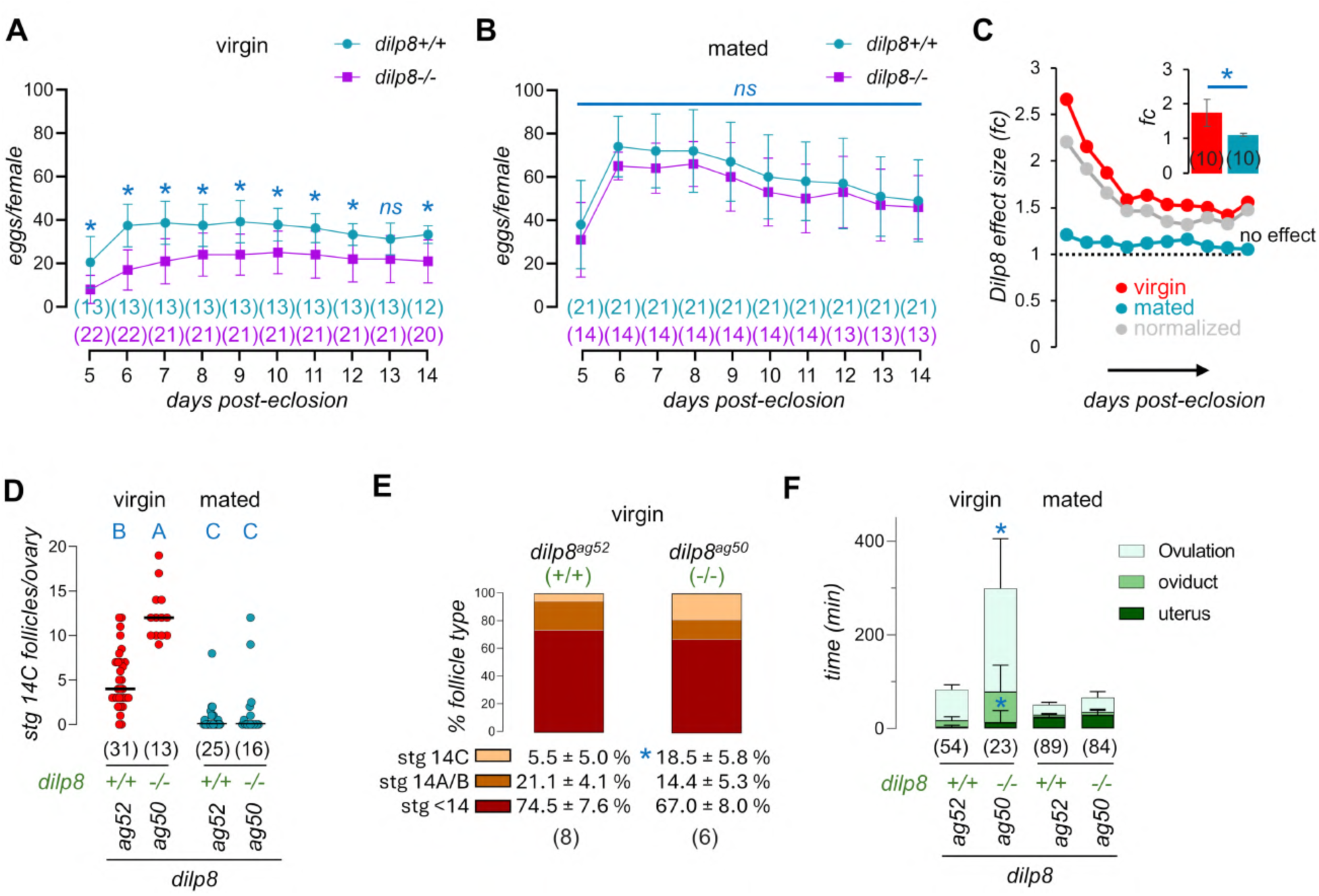
Dilp8 regulates ovulation rate. **(A)** Unfertilized egg laying evaluated for ten days is constantly significantly lower in virgin *dilp8* mutant flies (*dilp8^-/-^ = dilp8^ag50^*, magenta squares) against background control (*dilp8^+/+^ = dilp8^ag52^*, blue circles). **(B)** The effect is lost after mating. Shown is average ± SD of (*N*) repeats with five females each. Multiple t-tests with Bonferroni’s correction for multiple comparison: (*) *P* < 0.05; *ns*, *P* > 0.05. **(C) Dilp8 effect size (fold-change, fc): effect size of** the reduction (*dilp8^+/+^ / dilp8^- /-^*) of unfertilized egg laying or egg laying for virgin (blue) and mated (red) *dilp8* mutants, respectively. The reduction is higher in the first three surveyed days (days 5-7 post-eclosion). Normalized fold reduction (virgin/mated), gray. Inset: bars, average fc ± SD of *N =* 10 days (parentheses). * *P* < 0.0001, two-tailed, unpaired Student’s t-test. **(D)** Number of fully mature follicles (stage 14C follicles) per ovary (dots, *N* of ovaries in parentheses) is increased in virgin females of the indicated genotypes. Same blue letters, *P* > 0.05, Kruskal Wallis and Dunn’s test. **(E)** Proportion of fully mature (stg14C), early-14 (stage 14A/B) and earlier stages (1 to 13) in the ovaries (*N* in parentheses) of flies of the indicated genotypes. (*) *P* = 0.017, Student’s t–test with Bonferroni correction. **(F)** Duration of individual ovulation stages in flies (*N*) of the indicated genotypes. Ovulation time and uterus transit time, but not oviduct transit time, increase in mutant flies when compared to controls of indicated genotypes. Z-score test (*P* < 0.0005 and *P* < 0.005, respectively, both denoted as * in the graph). Duration of stages is not statistically-significant different between genotypes in mated females (*P* > 0.05, Z-score tests). See also Table S1.

Dissection of virgin female ovaries at day 6-7 showed that the number of fully mature stage (14C) follicles (Beard et al., 2023; Deady et al., 2017) in the ovaries of virgin *dilp8^ag50^* mutants was ∼3-fold higher than controls **(Fig. 3D).** This came at the cost of earlier stage 14 follicles (14A/B), but less so of follicles of other stages (**Fig. 3E**). These results suggested that control animals laid unfertilized eggs at a faster rate than *dilp8* mutants, so stage 14 follicles did not remain long enough in control ovaries to accumulate. In other words, ovulation is slower in *dilp8* mutants.

Quantification of ovulation and genital tract transit time using an established method (Beard et al., 2023) showed that ovulation took on average ∼80 min in control virgins, and eggs spent only ∼20% of that time in the oviduct and uterus. Mating reduced the overall ovulation time by ∼20% (to ∼65 min) but profoundly increased the fraction of time spent in transit in the oviducts (to ∼30 min or ∼50% of total ovulation time) (**Fig. 3F, Table S1**). In contrast to control virgins, *dilp8^ag50^* mutant virgins ovulated on average once every ∼300 min, but their oocytes spent a similar proportion of time in each compartment of the reproductive apparatus as controls (80% of the time in the ovary and 20% in the oviduct and uterus) (**Fig. 3F, Table S1**). Clearly, the whole ovulation process, rather than a specific phase, seems to be slower in *dilp8* virgins. In mated *dilp8* mutants, however, although ovulation rate was also slightly slower, residency times were not statistically significantly different from those in control mated females (**Fig. 3F**, **Table S1**). These results strongly suggest that virgin females use Dilp8 signaling for efficient ovulation, transit, and extrusion of unfertilized eggs from the uterus.

### The Dilp8 receptor, Lgr3, is required non-autocrinally for ovulation and egg laying in virgins

Virgin females homozygous for the *Lgr3^ag1^* null allele (Garelli et al., 2015), and hence unable to sense follicle-cell-derived Dilp8, also laid fewer unfertilized eggs than their respective background control (*Lgr3^ag2^*), a deficit that was rescued by mating (**Fig. 4A**). Using *Lgr3-GAL4* lines to infer *Lgr3* expression, a previous report suggested that *Lgr3* is expressed at the posterior tip of the stage 14 follicles (Liao and Nässel 2020), a region that overlays the aeropyle. Here, using a translational *Lgr3* reporter (*sfGFP::Lgr3^ag5^*) (Garelli et al., 2015), we found no detectable expression in this region. Instead, we found similar patterns of autofluorescence in the chorionic layers of the aeropyle in controls lacking any transgene (**Figs 1A, S3**). Consistently, analyses of single-cell RNA sequencing data does not reveal any detectable *Lgr3* expression in follicle cells (Li et al., 2022) (**Fig. S4)**. Namely, *Lgr3* is not detectable in female reproductive system (10x-crosstissue) posterior follicle cells, defined as *dilp8*+, *(pointed) pnt*+, and *(Matrix metalloproteinase 2) Mmp2*+ cells [the latter two, based on Zeng et al., 2024], or in any putative ovarian somatic cell, marked as negative for the germline marker, *vasa* (**Fig. S4)**. To further test for a possible autocrine role of Dilp8 on oocyte maturation, we dissected ovaries from virgin *dilp8* mutants and controls and performed qRT-PCR to quantify the expression of a subset of genes enriched in mature stage 14 follicles (*Oamb, CG15279, CG1698*, *CG3759,* and *CG6508*) (**Table S2**) (Tootle et al., 2011). Expression levels of these genes remained unchanged in *dilp8* mutants, indicating no consistent transcriptional response to the loss of Dilp8 (**Fig. S5)**. These results further support the hypothesis that Dilp8 does not act on follicle cells in an autocrine manner, but likely exerts its effect via other, *Lgr3*-positive cells.

**Fig. 4.**
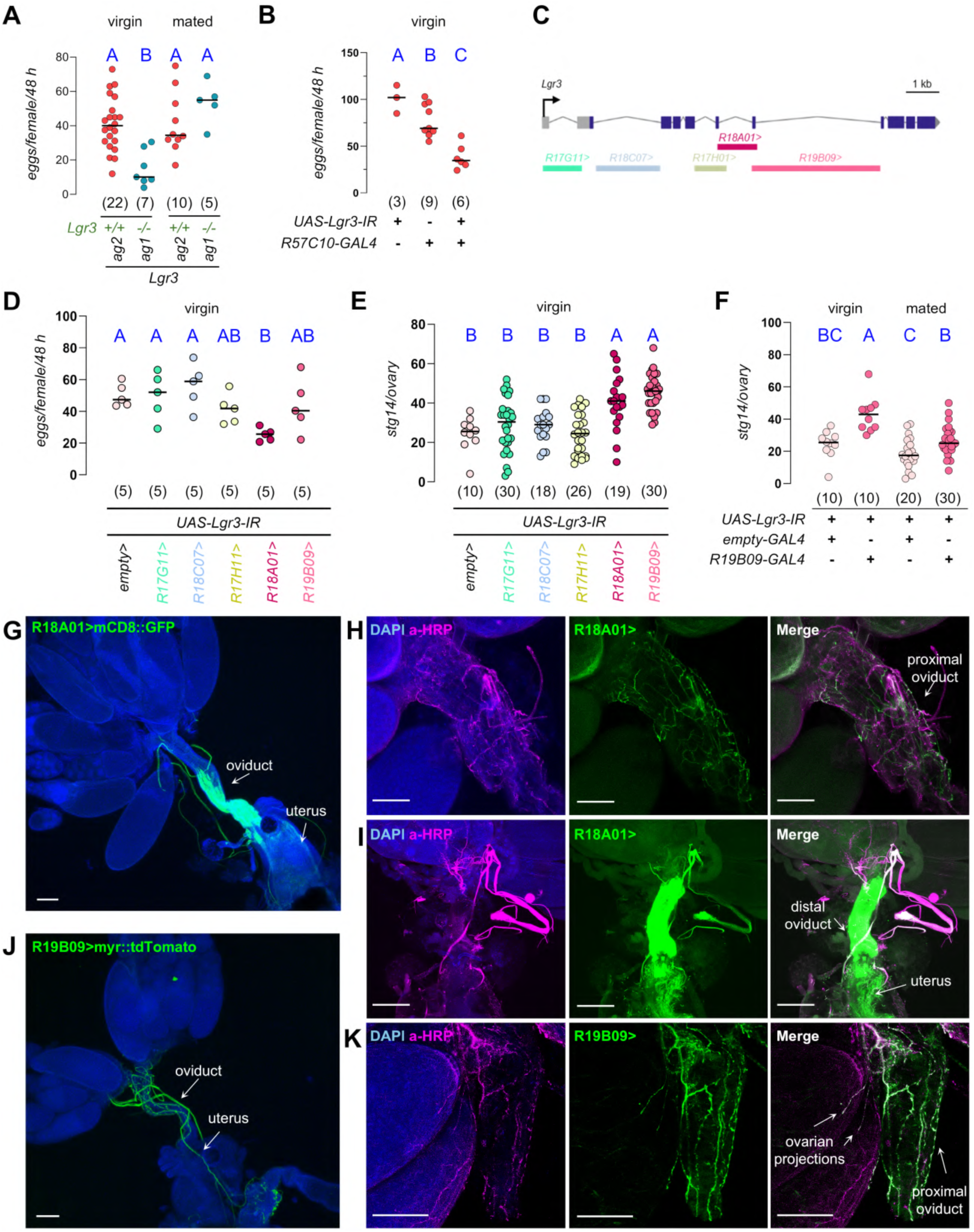
Lgr3 is required for virgin ovulation and egg laying in at least two different neuronal populations. **(A)** Virgin females lacking Lgr3 (*Lgr3^ag1^*), laid fewer eggs than their background controls (*Lgr3^ag2^*). The effect was lost upon mating. **(B)** Panneuronal downregulation of *Lgr3* by RNAi reduces the number of eggs laid by virgin females. **(C)** Schematic representation of the *Lgr3* locus depicting the regions spanned by the different *cis* regulatory modules that drive GAL4 expression. **(D-E)** Effect of the knockdown of *Lgr3* expression with individual *Lgr3 cis-regulatory module GAL4* lines on the **(D)** number of eggs laid and **(E)** number of mature oocytes stored in the ovaries of virgin females. **(F)** *Lgr3 RNAi* with *R19B09-GAL4* increases the number of mature follicles in the ovary of virgin females, and less so in mated females. Virgin data, same as in (E). *N* of (A,B,D) biological replicates, each with five individuals, and (E,F) ovaries in parentheses. Same blue letter, *P* > 0.05, ANOVA and Tukey’s HSD test. **(G-K)** Confocal Z-stack projections of the reproductive organs of females of the indicated genotypes. Fluorescent reporter expression is false colored in green. Nuclei labeled with DAPI (blue) and neuronal projections with anti-HRP antibody (magenta). **(H)** *R18A01>mCD8::GFP* marks neurons innervating the proximal basis of the oviduct. **(I)** Expression of *R18A01>mCD8::GFP* in the epithelial cells of the distal portion of the oviduct, and in neurons that project to the lower reproductive tract. **(K)** Expression of *R19B09>myR::tdTomato* in neurons that innervate oviduct and ovarian base. Scale bar = 100 µm.

### Lgr3 is required in different neuronal populations for ovulation and egg laying in virgins

A role for *Lgr3* on female reproductive physiology and metabolism has been proposed based on the finding that subsets of *Lgr3*-positive neurons affect female fecundity (Meissner et al., 2016) and sugar appetite (Laturney et al., 2023), and that an *Lgr3-GAL4* line can innervate the proximal oviduct (Liao and Nässel, 2020). It is unclear, however, if *Lgr3* is required for these neuronal activities. *Lgr3* transcripts are readily detectable in adult neurons by sequencing (Li et al., 2022) and by *in situ* hybridization (Meissner et al., 2016). Consistently, immunostaining shows sfGFP::Lgr3 protein in the adult nervous system of virgin and mated females, specifically in the brain and in the abdominal ganglion of the VNC (**Fig. S6**), from where many neurons innervate the female reproductive tract (Afkhami, 2024). By combining an RNAi line against *Lgr3* with the panneuronal driver *R57C10-GAL4*, we confirmed that *Lgr3* is required in neurons for proper laying of unfertilized eggs in virgin females (**Fig. 4B**). This requirement is coherent with a non-autocrine mechanism of follicle-cell-derived Dilp8 on the control of ovulation in virgins.

To narrow down the neuronal *Lgr3* requirement for ovulation control, we used a well-defined panel of five *cis-regulatory modules* from the *Lgr3* locus (Colombani et al., 2015; Garelli et al., 2015; Jaszczak et al., 2016; Vallejo et al., 2015) (**Fig. 4C**) to knock down its expression, and quantified unfertilized egg laying and the amount of stage 14 follicles in the ovary of virgins. Knockdown of *Lgr3* in *R18A01-GAL4* (*R18A01>*) cells significantly affected the number of unfertilized eggs laid by virgins whereas knockdown in *R17H01>* and *R19B09>* cells had a marginal or variable effect (**Figs 4D, S7**). Conversely, *Lgr3* knockdown in *R18A01>* and *R19B09>* cells led to a significant increase in the number of mature oocytes in the ovaries of virgins (**Fig. 4E, S7**). The *R19B09>Lgr3-IR* effect on oocyte retention was reduced by mating (**Fig. 4F**). Importantly, proper ovulation does not require *Lgr3* in *R17H01>* neurons, which had been shown to regulate fecundity when experimentally activated (Meissner et al., 2016). *Lgr3* is thus required in *R18A01>-*positive cells to control unfertilized egg laying and virgin ovulation and in *R19B09>-*positive cells to regulate virgin ovulation.

### Neurons requiring Lgr3 to control ovulation and egg transit innervate the reproductive tract

We verified the projection pattern of each of the five *Lgr3* GAL4 lines in the female reproductive system by driving membrane-tagged reporters (**Figs 4G,J, S10**). Interestingly, all of these lines, including the ones which we did not find a role for *Lgr3* in virgin unfertilized egg laying or ovulation, drove different expression patterns in the reproductive system consistent neuronal projections (**Fig. S10**), suggesting either that *Lgr3* is not expressed in them, or that *Lgr3* is expressed but required for other functions.

To confirm that the two cell populations where *Lgr3* was critical for either virgin unfertilized egg laying or oocyte retention, *R18A01>* and *R19B09>*, were non-overlapping, we placed a *LexA* version of *R19B09* [*R19B09>>* (Pfeiffer et al., 2010), which drives expression in similar neurons as the *GAL4* version, *R19B09>* (**Fig. S8**)] together with *R18A01*>, and found no overlap in the female reproductive system (**Fig. S9**). However, this experiment was limited because *R18A01>* drove very weak expression in neurons, where *Lgr3* is required (**Fig. 4B**). Using a stock carrying two copies of *R18A01>* (*2xR18A01>,* hereafter *R18A01>*) revealed a similar, yet stronger expression pattern as the single copy version, allowing us to more clearly define expression in fine neuronal fibers innervating the proximal base of the oviduct (**Fig. 4G,H**) and in neurons innervating the uterus (**Fig. 4G,I**). Additionally, *R18A01>* showed strong expression in epithelial cells of the oviduct, particularly in the distal portion, and the internal uterus lining (**Figs 4G,I, S9**). However, this epithelial expression unlikely reflects endogenous *Lgr3* expression, as single-cell RNA sequencing data is unable to detect *Lgr3* expression in oviduct cells (Li et al., 2022; Thayer et al., 2024). In contrast, *R19B09>myR::tdTomato* was strongly expressed in neurons innervating the whole oviduct, starting from the base of the ovary, with projections climbing up the ovary, and projections going down towards the distal oviduct (**Fig. 4J,K**).

We concluded that *Lgr3* is required in at least two neuronal populations to control ovulation and genital tract egg transit in virgins in response to follicle-cell derived Dilp8. From the expression pattern and different activities of each group of cells, we favor the hypothesis that Lgr3 in *R19B09>* neurons controls ovulation and oviduct transit, whereas Lgr3 in *R18A01>* neurons controls ovulation, oviduct transit, and uterus extrusion, leading to a stronger unfertilized egg laying phenotype than *R19B09>*.

### Neuroanatomical and neurotransmitter characteristics of neurons requiring Lgr3 to control ovulation and egg transit

Two major neuronal populations from the central nervous system send efferent projections to innervate the female reproductive tract (reviewed in Afkhami, 2024). A cluster of eight to ten modulatory octopaminergic neurons of the abdominal ganglion innervate the base of the ovary, oviduct, and the uterus (Middleton et al., 2006; Pauls et al., 2018), whereas other glutamatergic neurons, which form classical neuromuscular-junctions, innervate the oviduct and uterus [*e.g.*, (Cury and Axel, 2023)]. In addition, different sensory neurons innervate the reproductive tract and send afferent projections back to the central nervous system (Afkhami, 2024; Häsemeyer et al., 2009; Yang et al., 2009). The fact that both *R19B09>* and *R18A01*> drivers are expressed in the abdominal ganglia (Jenett et al., 2012; Pfeiffer et al., 2008) (**Fig. S11**), suggests they reach the reproductive system via efferent fibers.

To test these hypotheses more directly, we co-stained ovaries from animals expressing fluorescent reporters under *R18A01>* or *R19B09>* control with anti-Tdc2, which labels tyraminergic/octopaminergic (hereafter, octopaminergic) neurons (Pech et al., 2013), or anti-DVGlut, which labels glutamatergic neurons (Mahr and Abele, 2006; see methods and **Fig. S12**). As expected, *R19B09>*-positive fibers innervating the ovary were anti-Tdc2–positive, and a subset of *R19B09>*-positive fibers innervating the oviduct also colocalized with anti-Tdc2 (**Fig. 5A**). Another distinct set of *R19B09>*-positive neurons innervating the oviduct was anti-DVGlut–positive (**Fig. 5B,B’**). Thus, *R19B09>* labels both octopaminergic neurons innervating the ovary and oviduct and glutamatergic neurons innervating the oviduct. No *R19B09>*-positive projections were detected in the uterus.

**Fig. 5.**
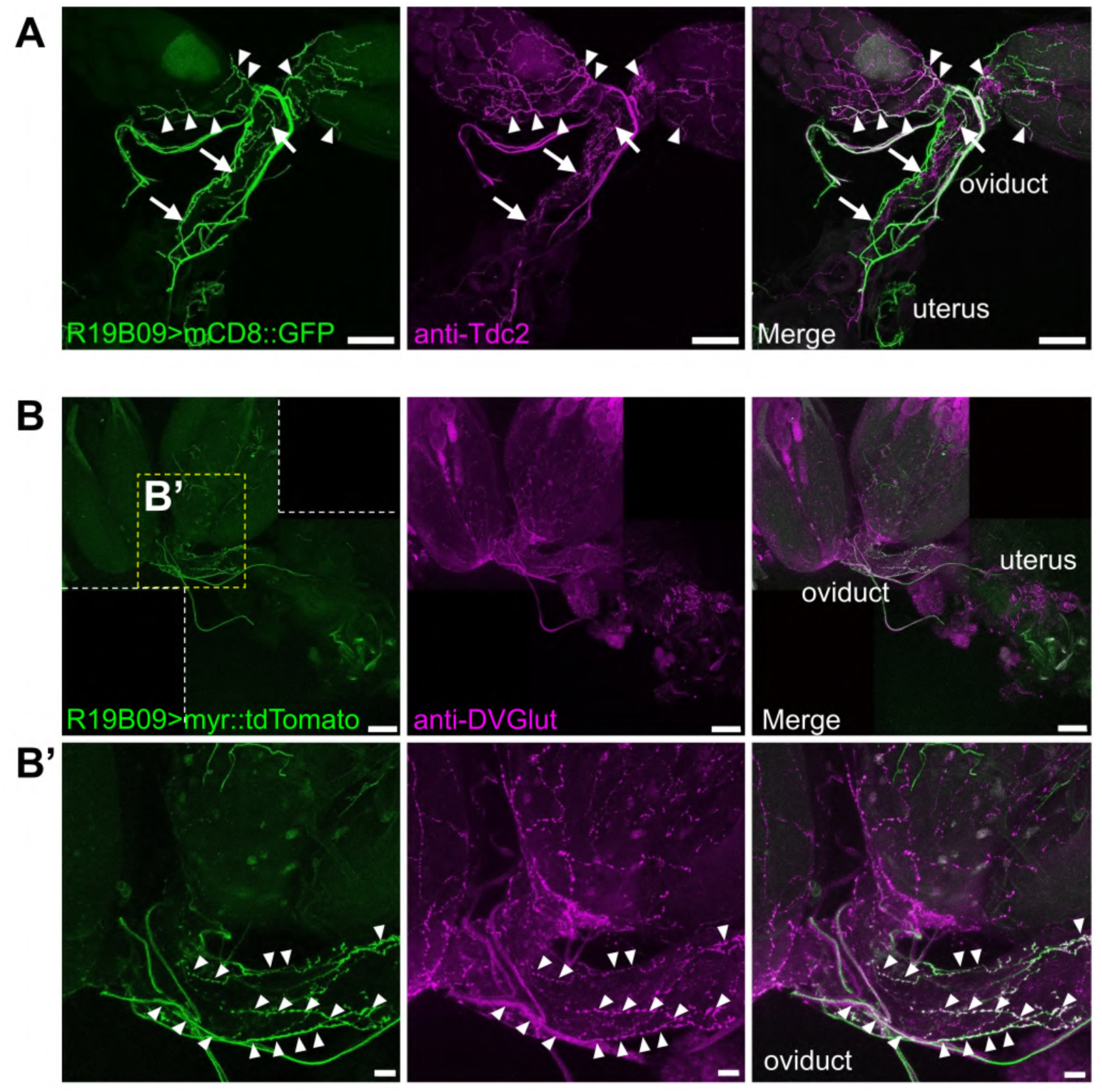
*R19B09>* is expressed in octopaminergic and glutamatergic neurons innervating the ovary and oviduct. **(A)** *R19B09>* is expressed in octopaminergic neurons that innervate the ovary (arrowheads) and oviduct (arrows). Montage of two projections of confocal z-stacks of a reproductive tract from a *R19B09>mCD8::GFP* mated female stained with with anti-GFP (green) and anti-Tdc2 (magenta). **(B, B’)** *R19B09>* is expressed in glutamatergic neurons innervating the oviduct. Projection of confocal z-stacks of a reproductive tract of a *R19B09>myr::tdTomato* (green) mated female stained with anti-DVGlut (magenta). **(B’)** Insert (yellow dotted line in (B)), depicting the proximal oviduct. Arrowheads depict neuronal fibers double positive for both markers. Scale bars: **(A)** 100 µm; **(B,B’,B’’)** 20 µm.

In contrast, *R18A01>*-positive neurons showed no detectable colocalization with anti-Tdc2 in either the oviduct or uterus (**Fig. S13**). Anti-DVGlut staining revealed uncertain overlap with a subfraction of thin *R18A01>*-positive fibers in the oviduct (**Fig. 6A,B**) and clear, substantial colocalization in the uterus wall, where terminals resembled neuromuscular junctions (**Fig. 6C**). Because the DVGlut-positive accumulations in the oviduct were difficult to assign definitively to *R18A01>*-positive fibers, we remain cautious about interpreting them as glutamatergic. Instead, most if not all of these thin lateral oviduct fibers may arise from peripheral sensory neurons known to innervate this region (*pickpocket-*positive (*ppk*+) cells; Häsemeyer et al., 2009; Lee et al., 2016; Yang et al., 2009), two of which are likely cholinergic (Rezaval et al., 2012). Collectively, these results indicate that at least a subset of *R18A01>*- and *R19B09>*-positive neurons are efferent, with *R18A01>* strongly marking glutamatergic fibers in the uterus and *R19B09>* marking both octopaminergic and glutamatergic oviduct-innervating fibers.

**Fig. 6.**
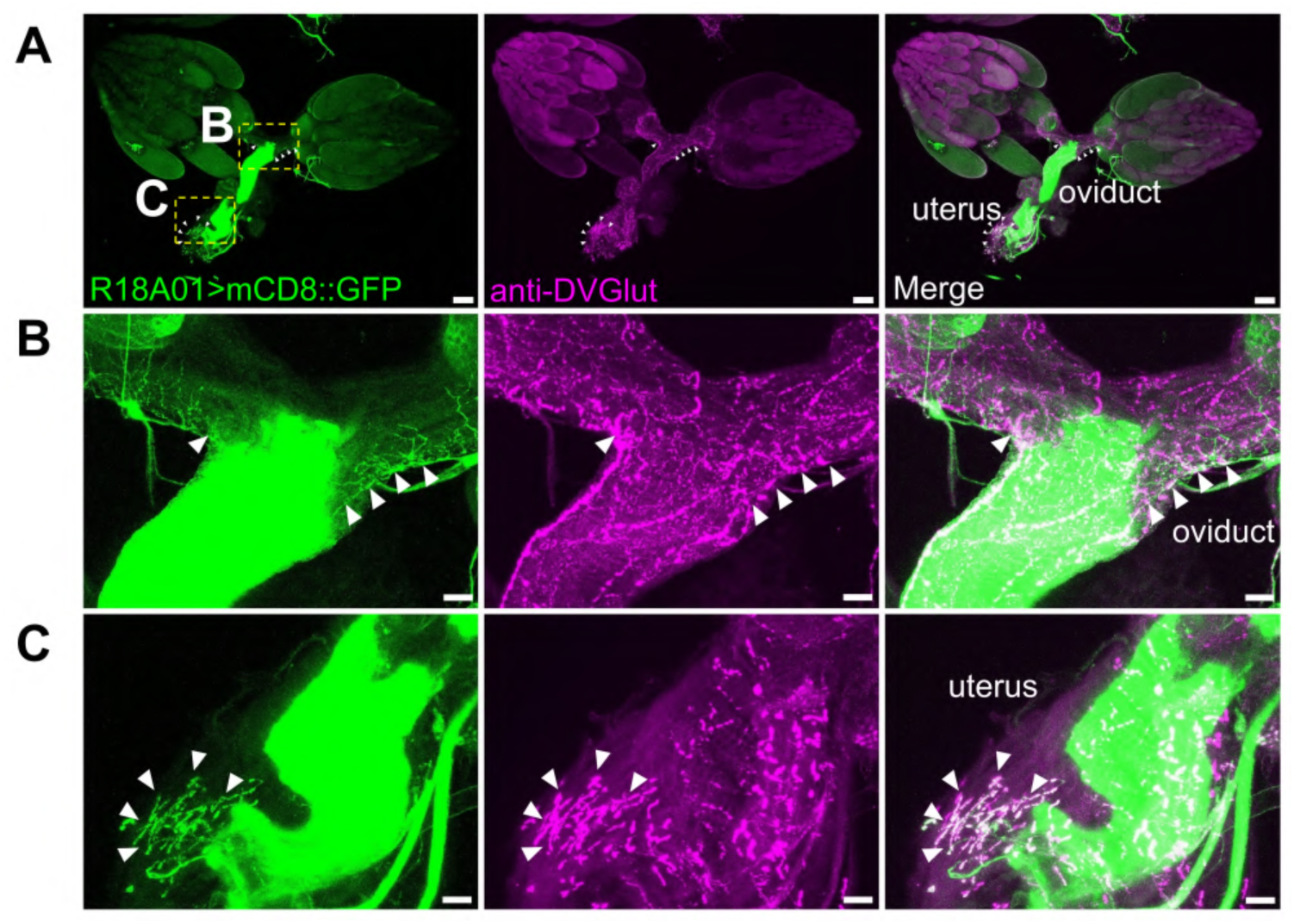
*R18A01>* is expressed in glutamatergic neurons innervating the oviduct and uterus. **(A)** *R18A01>* is expressed in glutamatergic neurons innervating the uterus and possibly the oviduct. Projection of confocal z-stacks of a reproductive tract of a *2xR18A01>mCD8::GFP* mated female stained with with anti-GFP (green) and anti-DVGlut (magenta). **(B)** Insert (yellow dotted line in (A)). Note fine neuronal fibers in the lateral oviducts. Arrowheads depict a subset of fibers that might be positive for both markers, although we favor the interpretation that both *R18A01>*-positive and anti-DVGlut-positive fibers are non-overlapping, but intimately associated in this region. **(C)** Same as (B) but focused on the inset around the uterine region in (A). Arrowheads, glutamatergic motorneuron synaptic termini at the outer uterine wall. Scale bars: (A) 100 µm; (B,C) 20 µm.

### Dilp8 indirectly promotes oocyte quality by facilitating ovulation of older oocytes

*Drosophila* oogenesis is a relatively energy-intensive process and unfertilized egg laying is considered a pointless loss of reproductive potential (Akhund-Zade et al., 2017; Boulétreau-Merle and Fouillet, 2002; Horváth and Kalinka, 2018). However, this must be contrasted with the observations that prolific unfertilized egg laying (*i.e.,* short retention) phenotypes are selected for in warmer climates and the potential for expressing such phenotypes remains present in cold-adapted wild populations via balancing selection (Boulétreau-Merle, 1990; Boulétreau-Merle et al., 1992; Boulétreau-Merle and Fouillet, 2002). Hence, it is likely that short ovulation times are subject to selective pressures, just like longer retention ones are subject to pressures for overwintering.

Similar to human oocytes (te Velde and Pearson, 2002) and those of *Caenorhabditis elegans* (Luo et al., 2010), older mature oocytes in *Drosophila* display reduced viability (hatchability), which degrades proportionally to the time the oocyte spends in retention in the ovary (Greenblatt and Spradling, 2018; Greenblatt et al., 2019). This alone could provide a selective force for the Dilp8-Lgr3 pathway to promote ovulation in virgins: to frequently purge the ovary of older, less viable oocytes. To evaluate the effect of egg retention on viability and any additional role Dilp8 could have in promoting oocyte quality, we assayed hatchability in *dilp8* mutant and control flies.

For this, we grew well-fed *dilp8-eGFP^MI00727^* or *dilp8^ag50^* mutants and their respective controls (*w^1118^* or *dilp8^ag52^*) for 4 days on yeast-enriched food and then switched them to hardened agar apple-juice plates for 7 days. This regimen drastically reduces virgin egg-laying, leading to mature oocyte aging during the retention period. On day 11, these flies were mated, and we scored the % of eggs that hatched into larvae as a readout of oocyte quality (**Fig. 7A**). We found that both controls that retained oocytes for 7 days as virgins had a drastic reduction in hatchability compared to their continuously mated and fed counterparts (**Fig. 7B,C**). However, the difference was no longer observed two days after mating (day 13), indicating that the loss of oocyte quality is due to the prolonged retention of the older oocytes in the ovary (**Fig. 7B,C**). The same drastic reductions in hatchability at day 11 and recoveries at day 13 occurred in *dilp8-eGFP^MI00727^*and *dilp8^ag50^* mutants that retained oocytes, and these reduction and recoveries were not statistically significantly different from those of controls (**Fig. 7B,C**). The conclusions of these experiments are two-fold: first, these results clearly show that old oocytes have reduced hatchability in independent genetic backgrounds, consistent with previous reports (Greenblatt and Spradling, 2018; Greenblatt et al., 2019), providing a suitable mechanistical explanation for the role of the Dilp8-Lgr3 pathway in virgin females: to promotes oocyte quality by facilitating–via ovulation–the elimination of older oocytes. Second, we could not reject the null hypotheses that Dilp8 does not promote oocyte quality by an additional mechanism, apart from by promoting ovulation in virgins.

**Fig 7.**
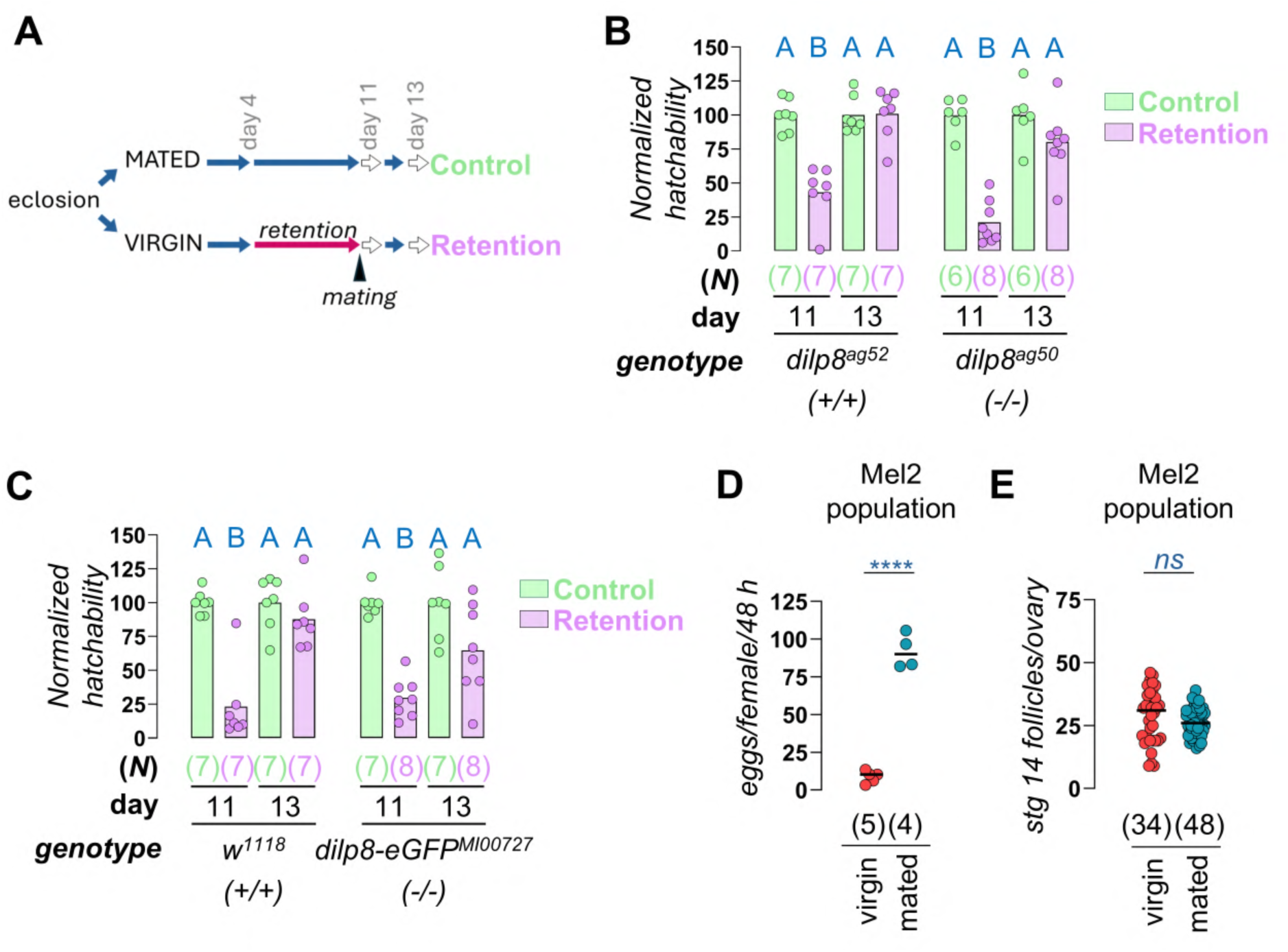
Egg retention leads to reduced hatchability. **(A)** Protocol used to induce egg retention. Hatchability was evaluated on the eggs laid at days 11 and 13 after eclosion. Eggs laid at day 13 after eclosion are considered newly generated and non-retained. See details in Methods section. **(B-C)** Egg viability is reduced after seven days of oocyte retention (day 11, pink) in the ovary of different *dilp8* mutant flies and their background controls according to the depicted genotypes. Hatchability of newly generated oocytes is restored to control values (green bars) two days later (day 13). Each dot represents an independent biological replicate (*N*) with five flies each (bar, average). **(D)** Egg laying and **(E)** number of stage 14 follicles per ovary in virgin and mated females from an outbred *D. melanogaster* population. ****, *P* < 0.0001, ns, *P >* 0.05, unpaired two-tailed Student’s t-test. (B,C,D) *N* of replicates, each with five individuals and (E) *N* of ovaries in parentheses. (B,C) Same blue letters, *P* > 0.05, ANOVA and Tukey’s HSD test.

### dilp8 mutants are in a short-retention background: implications for oogenesis and oocyte quality control

Whereas using laboratory stocks, such as *w^1118^* “controls”, allows us to better control for genetic backgrounds, it is not clear how faithfully they reflect the behavior of wild fly populations in our experimental settings. To get a better perspective of this, we profiled virgin and mated egg-laying behavior in a previously-characterized outbred *D. melanogaster* population (Mel2) from Portugal (temperate climate) (Faria and Sucena, 2017; Martins et al., 2013). Our results showed that well-fed virgins spontaneously laid 4.3 ± 2.0 (average ± SD) unfertilized eggs per day (∼1 ovulation every 5-6 h), which rose to 45.9 ± 5.7 eggs per day (1 ovulation every 0.5 h) after mating (**Fig. 7D)**. This result has many implications. First, it confirms that spontaneous ovulation is neither a negligible process in the wild nor an artifact of stocks kept under laboratory conditions for decades. Besides, it is in line with previous reports of other flies recently collected from the wild or with their established populations that displayed virgin egg laying (Akhund-Zade et al., 2017; Boulétreau-Merle and Fouillet, 2002; Horváth and Kalinka 2018;). Second, it shows that while Mel2 population virgins align more with the long-retention phenotype, the virgins of the various laboratory controls we profiled herein that lay 20-30 unfertilized eggs per day (**Fig. 2B,C,D, 3A, 4A,B,D**) are more representative of short retention types (Boulétreau-Merle, 1990; Boulétreau-Merle and Fouillet, 2002). This genetic background serendipitously facilitated the detection of a role for the Dilp8-Lgr3 pathway in promoting ovulation and egg transit through the genital tract. Third, virgin and mated females of the Mel2 population accumulated stage 14 oocytes in their ovaries to a similar extent (**Fig. 7E)**. This indicates that despite laying even fewer unfertilized eggs per day than virgin *dilp8* mutants (which lay ∼5-10 unfertilized eggs per day), virgins of the Mel2 population do not accumulate more than 1-2 stage 14 follicles per ovariole (considering ∼20 ovarioles per ovary [as a reference, the DGRP lines have 19.4 ± 2.84 average ± SD ovarioles per ovary, *N =* 200 lines (Lobell et al., 2017)]). In fact, long retention virgin females, such as Mel2 population females, do not accumulate stage 14 follicles beyond 2 per ovariole because they abruptly reduce new follicle production (Boulétreau-Merle, 1990; Boulétreau-Merle and Fouillet, 2002). This suggests the existence of an oogenesis progression checkpoint that is sensitive to the amount of stage 14 follicles per ovariole.

In contrast, virgins with short retention phenotypes lay copious amounts of unfertilized eggs and do not reach the “2 mature oocyte/ovariole” steady state. This suggests that the checkpoint either does not exist in short-retention flies, or that it does exist but is never activated due to continuous unfertilized egg laying. In the latter scenario, virgin *dilp8* mutant females, which are in an apparent short-retention background (*w^1118^*), would accumulate mature oocytes beyond the threshold because checkpoint-mediated arrest of oogenesis requires Dilp8 activity. This interpretation would align with Dilp8 having a second independent role promoting oocyte quality: to antagonize oogenesis (more precisely, vitellogenesis) progression. This could, for instance, reduce the generation of mature oocytes that would age for prolonged periods in the ovary. If this were true, one would expect a slower recovery of hatchability in *dilp8* mutant animals that retained oocytes, which we do see in both genetic backgrounds, but the reduction is not statistically significant (compare two last columns in **Fig. 7B,C**).

The negative feedback by which long-retention animals halt new mature stage 14 follicle production kicks in after they accumulate ∼2 stage 14 follicles per ovary (Boulétreau-Merle. 1990; Boulétreau-Merle and Fouillet, 2002), strongly suggesting that it responds to the amount of stage 14 follicles, and making Dilp8 secreted from follicle cells a candidate molecule signaling the negative feedback. This negative feedback pathway is reminiscent of the nutrition-dependent checkpoint, which prevents development of vitellogenic oocytes beyond stage 8/9 in the absence of protein-rich food sources (Drummond-Barbosa and Spradling, 2001; Greenblatt and Spradling, 2018; Greenblatt et al., 2019; Richard et al., 2005) and the gating mechanism that limits egg production while conditions are suboptimal in stressful conditions (Meiselman et al., 2017; Meiselman et al., 2018). These likely correspond to the same checkpoint, which is controlled at multiple levels, including by mating (Soller et al., 1999).

Indeed, oogenesis progression is subject to extensive endocrine control by the insulin/insulin-like growth factor (hereafter, insulin), ecdysone, ecdysis triggering hormone, and juvenile hormone signaling pathways (Drummond-Barbosa and Spradling 2001; Greenblatt and Spradling 2018; Greenblatt et al., 2019; Gruntenko et al., 2010; Liu et al., 2008; Meiselman et al., 2017; Meiselman et al., 2018; Richard et al., 2005; Soller et al., 1999;). Here, we hypothesized that the Dilp8-Lgr3 pathway, beyond promoting ovulation and egg transit through the reproductive tract, also regulates the oogenesis progression checkpoint by negatively affecting insulin signaling. This would mean that insulin signaling would be somehow increased in *dilp8* mutants. We tested this by quantifying an insulin signaling readout (Akt phosphorylation) and *dilp3* mRNA levels, which had been previously antagonistically linked to Dilp8 (Li et al., 2023; Liao and Nässel, 2020), in different *dilp8* and *Lgr3* mutants, but found no robust or consistent effect (**Figs S14, S15**). The mechanism by which Dilp8-Lgr3 antagonize oogenesis progression remains to be defined.

## DISCUSSION

### The Dilp8-Lgr3 pathway promotes ovulation and unfertilized egg transit through the genital tract in virgins

A role for Dilp8 in controlling ovulation, fecundity, and metabolism in mated females had been previously proposed based on RNAi studies and a hypomorphic mutation (Li et al., 2023; Liao and Nässel, 2020). Here, we show that the Dilp8-Lgr3 relaxin-like pathway indeed modulates ovulation and genital tract egg transit, but also oocyte quality, and it does so most clearly and strongly in virgin females. We show that these roles are less visible and consistent in mated animals, suggesting that mating provides redundant inputs that largely override the requirement of the Dilp8-Lgr3 pathway.

Dilp8 is upregulated in fully mature, stage 14 follicles, where it was proposed to act autocrinally on cells at the posterior tip, based on the expression of fluorescent reporters driven by Lgr3 regulatory regions (Liao and Nässel, 2020). In contrast, we show that this region is strongly autofluorescent due to the underlying chorionic structure, called aeropyle, and find no detectable expression of *Lgr3* with an extensively used Lgr3 protein reporter construct (Fernandez-Acosta et al., 2025; Garelli et al., 2015; Heredia et al., 2021). Also arguing against an autocrine role in follicle maturation is the lack of effect of *dilp8* mutation on oocyte maturation markers and the observation that the egg laying deficit of virgin *dilp8* mutants is largely rescued by mating, indicating that ovarian dynamics can proceed normally without *dilp8* under these conditions. Altogether, these results suggest that Dilp8 acts in a paracrine manner in nearby tissues and/or in a systemically rather than on follicle cells themselves.

A conceptually important finding is that Dilp8 is secreted from stage 14 follicle cells in an ovulation-independent manner when they still reside in the ovary, likely basally/basolaterally into the ovariole/oviduct space, from where it could reach target tissues. Functional assays revealed that Dilp8 acted in at least two different *Lgr3+* populations of neurons marked by *R18A01* and *R19B09 cis-regulatory modules* with projections to complementary regions of the reproductive tract. While the simplest explanation is that Dilp8 acts on these subpopulations of neurons, we cannot exclude the involvement of other neurons that do not innervate the reproductive tract. Despite this limitation, our results suggest a new model where follicle-cell-derived Dilp8 acts either as a paracrine or systemic signal on multiple Lgr3+ neurons (**Fig. 8**).

**Fig. 8.**
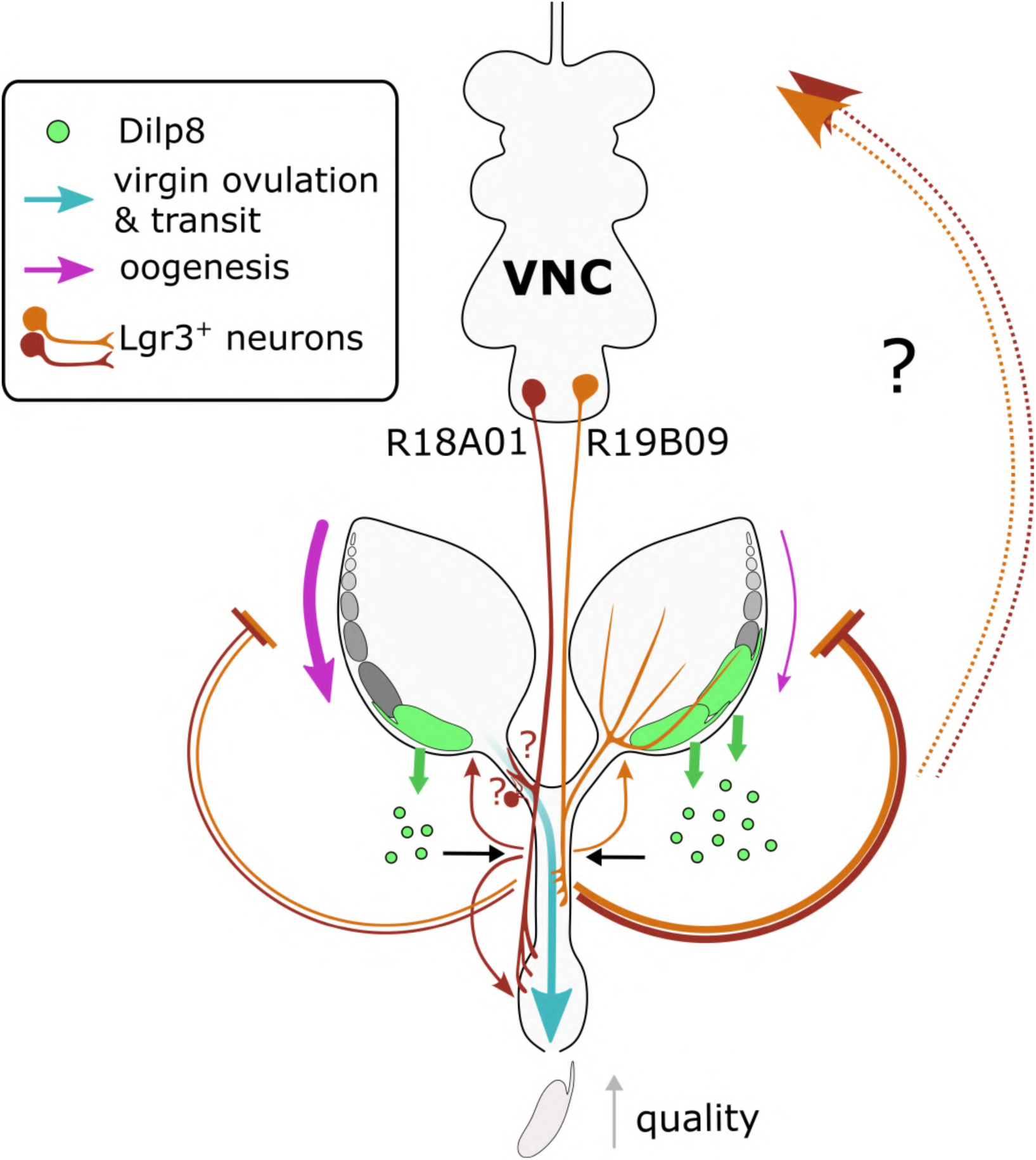
Model for the role of the Dilp8-Lgr3 signaling pathway in virgin reproductive physiology control. Dilp8 peptide (green circles) continuously secreted from fully mature Dilp8+ follicles (green follicles), acts (black arrows) on two populations of Lgr3-expressing neurons (brown and light brown neurons) that innervate the lateral oviduct and uterus (R18A01), and the ovary and the lateral and common oviducts (R19B09). Activities from these neurons (thin curved arrows) facilitate ovulation and egg transit through the reproductive tract (large blue arrow) and inhibit oogenesis progression (magenta arrow) in virgin females, contributing to keeping the number of fully mature follicles below 2 per ovariole in the ovaries. In this model, the soma of both *R18A01* and *R19B09* neurons reside in the abdominal ganglion of the ventral nerve cord (VNC), but this remains to be fully confirmed (black question mark). It is possible that the lateral oviduct-innervating *R18A01* neurons are sensory neurons (brown neuron near question mark). Collectively, these activities promote egg quality.

### Characteristics of the Lgr3+ neurons mediating Dilp8 activity on female virgin reproductive physiology

We found that ovary-innervating fibers of *R19B09>* neurons are octopaminergic, as expected, and so are a few ramifications descending the oviduct. The rest of the *R19B09>-*positive oviduct projections are clearly glutamatergic. Although these expression patterns align with both the function we attribute to Lgr3 in *R19B09>* neurons—promoting ovulation in virgins—and their presumed origin as efferent abdominal ganglion projections, it remains to be defined which neuronal subpopulations require Lgr3 to promote ovulation and how these neurons act.

*R18A01>* drives expression in glutamatergic neurons innervating the uterus with a neuroanatomy consistent with classical neuromuscular synapses (Daniels et al., 2004; Jan and Jan, 1976), in line with the detection of Lgr3 protein- and *R18A01>*-positive cells in the VNC abdominal ganglion in this study (**Fig. S6 and Fig. S11**) and *Lgr3* transcripts and *Lgr3-GAL4*-positive expression in other studies (Liao and Nässel 2020; Meissner et al., 2016). These neurons resemble the glutamatergic circular muscle of the uterus (CMU) neurons, which promote egg laying (Cury and Axel, 2023). Future work could test if CMU neurons are Lgr3+ and respond to follicle-derived Dilp8, predictably facilitating CMU neuron activation and egg extrusion.

*R18A01>* is also expressed in the proximal oviduct as very thin fibers that resemble cholinergic *ppk*+ sensory neurons (Häsemeyer et al., 2009; Rezaval et al., 2012; Yang et al., 2009) *ppk+* neurons play critical roles in female reproductive behavior in the sex peptide pathway (Häsemeyer et al., 2009; Yang et al., 2009; Yapici et al., 2008). Silencing all *ppk+* neurons in virgins, but not mated animals, induces robust egg laying (Häsemeyer et al., 2009; Yang et al., 2009), a phenotype reminiscent of short retention animals, which we show here requires a functional Dilp8/Lgr3 pathway. If these *R18A01* neurons are indeed *ppk+*, it would be interesting to investigate interactions between these pathways. Independently, the expression patterns described above are consistent with the functions we ascribe for *Lgr3* in *R18A01>* neurons: ovulation and unfertilized egg laying promotion in virgin females. The definition of which of these neurons actually express Lgr3 protein and are responsible for each activity will require further work.

### The Dilp8-Lgr3 pathway ensures optimal oocyte quality by two possible mechanisms

The promotion of spontaneous ovulation by the Dilp8-Lgr3 pathway in virgin females increases average oocyte quality by discharging older oocytes, which have reduced hatchability rates. Considering that *dilp8* mutant virgins accumulate more than 2 mature oocytes per ovariole, this suggests a compromised checkpoint for regulating oogenesis progression, which could generate lower quality oocytes by a second mechanism (**Fig. 8**). A well-characterized vitellogenesis checkpoint is controlled by systemic insulin signaling and is sensitive to nutritional and stress conditions (Drummond-Barbosa and Spradling, 2001). However, we detected no consistent effects of Dilp8 or Lgr3 activity on insulin-signalling-dependent Akt phosphorylation in the female body or on *dilp3* mRNA levels, which had previously been suggested to be modulated by Dilp8 (Li et al., 2023; Liao and Nässel, 2020) across different genetic mutations and backgrounds (**Figs. S14, S15**). These findings could mean that Dilp8 antagonizes insulin signaling only in certain genetic backgrounds, or, alternatively, that the Dilp8–Lgr3 pathway is not directly connected to insulin pathway regulation in virgin females. The apparent genetic interactions may instead reflect uncontrolled factors such as cryptic background mutations or nutritional variation, including microbiome differences. Further work is clearly needed to understand why this checkpoint fails when Dilp8 or Lgr3 function is reduced.

A similar phenotype is observed in mutants partially deficient in the octopamine pathway, which show a strong accumulation of more than two stage-14 oocytes per ovariole primarily due to ovulation defects (Cole et al., 2005; Middleton et al., 2006; Monastirioti et al., 1996; Monastirioti, 2003). It will be informative to determine whether their oocyte accumulation can be uncoupled from the ovulation defect, and whether the Dilp8–Lgr3 pathway remains functional in these backgrounds. Dual control of ovulation and oogenesis by Dilp8–Lgr3 also resembles the ecdysis-triggering hormone pathway, which regulates ovulation via octopamine and oogenesis via juvenile hormone (Meiselman et al., 2018). INKA-cell–ablated females, lacking detectable *ecdysis-triggering hormone* mRNA, accumulate late-stage oocytes due to ovulation defects but not more than 2 mature oocytes per ovariole, even when ectopic juvenile hormone is provided (Meiselman et al., 2018). This suggests that the checkpoint remains active and that Dilp8–Lgr3 and ecdysis-triggering hormone pathways likely act through distinct yet complementary mechanisms.

Two recent studies examined Dilp8–Lgr3 signaling in ovulation. Oramas et al. (2025) largely support our findings, confirming a role for this pathway in spontaneous ovulation in virgins, though disparities—such as their stronger R19B09> neuronal Lgr3 knockdown phenotype—require clarification. In contrast, Yang et al. (2025) report Lgr3 acting in octopaminergic neurons after mating and find no role in virgin ovulation. Differences in assays (24-48-h long-term egg-laying vs. 3-h ovulation snapshots) and genetic tools (null mutations and VALIUM22 vs. weaker VALIUM10 RNAi; Garelli et al., 2015) likely explain these conflicts, suggesting the results may ultimately be reconcilable.

### Conserved roles of the Dilp8-Lgr3 pathway in female reproductive physiology

The relaxin pathway (Bathgate et al., 2013) has been historically associated with female reproduction since it was first demonstrated to promote relaxation of the interpubic ligament in pregnant guinea pigs and rabbits (Hisaw, 1926) and to be produced by the corpus luteum (Fevold et al., 1930). It has become clear that the relaxin pathway was present in the last bilaterian ancestor (Feng et al., 2025; Gontijo and Garelli, 2018; Mita et al., 2009), and that it has evolved multiple functions that go beyond female reproduction, including functions on tissue remodelling, multi-tissue physiology regulation, behavior, stress regulation, and development (Bathgate et al., 2013; Gontijo and Garelli, 2018; Heredia et al., 2021; Imambocus et al., 2022), making relaxins potential therapeutic targets for multiple diseases (Bathgate et al., 2018). Although relaxins have mostly been associated with late pregnancy events in vertebrates [reviewed in (Klein, 2016)], relaxin is also expressed in follicles and other ovarian cells in many vertebrates, including during the nonpregnant oestrus cycles, and a direct role in promoting ovulation is clear in horses and rats, where relaxin induces the secretion of enzymes that stimulate follicle cell rupture [reviewed in (Klein, 2016)]. A role for relaxin signaling in ovulation/spawning control was also recently demonstrated in starfish (Feng et al., 2025). Our study in *Drosophila*, a protostome, therefore allows for the conclusion that the modulation of spontaneous ovulation is an ancestral role of the relaxin pathway in bilaterians.

## MATERIALS AND METHODS

### *Drosophila* husbandry and stocks

The following stocks were used in this study:

*w*; P{UAS-shi^ts^1.K}3* was a gift from Rita Teodoro. The *Lgr3^ag1^* mutant, the *sfGFP::Lgr3^ag^*^5^ reporter, and *Lgr3^ag2^* and *Lgr3^ag11^* background controls were previously described (Heredia et al. 2015). The *dilp8^ag50^*and *dilp8^ag54^* mutant alleles and their background control *dilp8^ag52^* were previously described (Heredia et al. 2021). *P{UAS-dpp.sGFP} (UAS-sGFP)* (Entchev et al., 2000) and *dilp8^KO^* (Boone et al. 2016) were gifts from Pierre Leopold. *empty-GAL4* is *pBDP[GAL4] attp2 (III)* and was a gift from Carlos Ribeiro. Our *w^1118^* stock can be tracked back to the Dominguez lab, Alicante, Spain. *UAS-Lgr3-IR* is a TRiP VALIUM22 short hairpin insertion derived from the stock *y^1^ sc* v^1^ sev^21^; P{y^+t7.7^ v^+t1.8^=TRiP.GL01056}attP2* (BL36887) (Ni et al., 2011), obtained from the Bloomington Drosophila Stock Center, Indiana University Bloomington (BDSC; BL, stock number), and then placed in a *w^1118^* background and kept balanced over *TM6B*. *dilp8-1-GAL4* is *y* w*; P{GawB}NP0007* and *dilp8-2-GAL4* is *w*; P{w^+mW^.hs=GawB}NP0778*, and both were obtained from the Kyoto Drosophila Stock Center. *dilp8-1-GAL4* was recombined with *w*; P{y^+t7.7^ w^+mC^=10XUAS-IVS-mCD8::RFP}attP40* (BL32219) and then backcrossed to *w^1118^* to get *w^1118^; dilp8-1>mCD8:RFP* (*dilp8-1>*), as described below. *y^1^ w*; Mi{MIC}Ilp8^MI00727^* (BL33079, *dilp8^MI00727^*) was backcrossed to *w^1118^* to get *w^1118^; dilp8^MI00727^*, as described below. *UAS-dilp8-IR* is a recombinant of *y^1^ v^1^; P{y^+t7.7^ v^+t1.8^=TRiP.HMS06016}attP40* (BL36726) with *P{KK112161}VIE-260B (v102604)*, obtained from the Vienna Drosophila Resource Center, respectively. The following stocks were also obtained from the BDSC:

- *w^1118^; P{y^+t7.7^ w^+mC^ =GMR47A04-GAL4}attP2* (BL50286)
- *w^1118^; P{y^+t7.7^ w^+mC^=GMR57C10-GAL4}attP2* (BL39171)
- *w^1118^; P{y^+t7.7^ w^+mC^=GMR19B09-GAL4}attP2* (BL48840)
- *w^1118^; P{y^+t7.7^ w^+mC^ =GMR17G11-GAL4}attP2* (BL49275)
- *w^1118^; P{y^+t7.7^ w^+mC^=GMR18C07-GAL4}attP2* (BL48806)
- *w^1118^; P{y^+t7.7^ w^+mC^=GMR18A01-GAL4}attP2* (BL48791)
- *w^1118^; P{y^+t7.7^ w^+mC^=GMR17H01-GAL4}attP2* (BL48786)
- *w^1118^; P{y^+t7.7^ w^+mC^=GMR19B09-lexA}attP40* (BL52539)
- *w*; P{y^+t7.7^ w^+mC^=10XUAS-IVS-myr::tdTomato}attP40* (BL32222)
- *w^1^; P{y^+t7.7^ w^+mC^=10XUAS-IVS-myr::GFP}attP40* (BL32198)
- *w*; P{y^+t7.7^ w^+mC^=20XUAS-IVS-mCD8::GFP}attP2* (BL32194)
- *y^1^ w* P{y^+t7.7^ w^+mC^=10XUAS-IVS-mCD8::RFP}attP18 P{y^+t7.7^ w^+mC^=13XLexAop2-mCD8::GFP}su(Hw)attP8* (BL32229)
- *y^1^ w*; P{w^+mC^=UAS-mCD8.mRFP.LG}10b* (BL27399)
- *y^1^ w^67c23^* (BL6599)
- *y^1^ v*;; P{CaryP}attP2* (BL36303)
- *w*; TI{RFP[3xP3.cUa]=TI}Lgr3[attP]/TM3, Sb^1^* (BL84516) (Deng et al., 2019)

Stocks are maintained at low densities at 18 °C in a 12-h light/dark cycle. All experiments were done at 25 °C in a 12-h light/dark cycle, or at a different temperature when specified.

Backcrossed stocks had their 1st and 2nd chromosomes substituted for those of the *w^1118^*stock using directional crossing and balancer chromosomes (CyO), while the 3rd chromosome was backcrossed 5x in total.

An outbred *D. melanogaster* population (Mel2) was obtained from the Sucena lab (Faria and Sucena 2017; Martins et al. 2013). The population is maintained in laboratory cages at a census of 1500-2000 individuals, under constant temperature (25°C) and humidity (60-70%) with a 12-h light/dark cycle. All experimental flies were generated from egg lays with controlled density.

Food: Animals are fed with cornmeal-agar medium, consisting of 4.5% molasses, 7.5% sugar, 7% cornflour, 2% granulated yeast extract, 1% agar, and 0.25% nipagin, mixed in distilled water.

### Precise excision of the *Mi{MIC}* element from *dilp8-eGFP^MI00727^*

The precise excision of the *Mi{MIC}* element from *dilp8-eGFP^MI00727^* (Venken et al., 2011), which we named *dilp8^ag22^*, was obtained by crossing virgin *w^1118^; dilp8-eGFP^MI00727^* backcrossed females with males carrying the heat shock-inducible MiET1 transposase on a *Cy* balancer chromosome (*w^1118^; If/Cy-Minos; MKRS/TM6B*) and transferred to new vials every day. After 48 h of development and until pupariation, the F1 progeny was given a 1-h 37 °C heat shock daily to induce the expression of the MiET1 transposase. *Cy-Minos/+; dilp8-eGFP^MI00727^/MKRS* or *TM6B* male adults were selected and individually crossed to the balancer strain *w^1118^; If/CyO; MKRS/TM6B*. Deletion of the *Mi{MIC}* element was confirmed by PCR with primers dilp8MMIC_F CAATAGATCAGTAGGCACGTCGAT and dilp8MMIC_R CAGCAAGTCCAGCAGCATGT and Sanger sequencing. *w^1118^; If/CyO; dilp8^ag22^/TM6B* animals were then crossed with *w^1118^; Chr II from w^1118^; dilp8^ag22^/TM6B*, generating *w^1118^;;dilp8^ag22^*. The restoration of wild type *dilp8* expression level was confirmed by qRT-PCR (**Fig. 1C**).

### Egg-laying assay and hatchability assessment

Flies used to evaluate oogenesis and oviposition were reared under controlled conditions. Females laid eggs overnight on apple juice egg-laying plates (25% apple juice, 2.5% sucrose, 2% agar, 5% nipagin in a 60-mm diameter petri dish). The next day, L1 larvae were transferred to vials with fly food at low density (∼30 larvae per vial) and kept at 25 °C. Virgin females reared under these conditions were placed in groups of ∼10-15 per vial with paste yeast supplemented food until they were used for egg-laying assays four-five days later. At that time, they were transferred to egg-laying plates supplemented with yeast paste, at a rate of five females per cage. Egg-laying plates were replaced every 24 h for a total of 48 h, except in long-term oviposition experiments, where flies were evaluated for 10 d.

The number of eggs laid on the plates was counted at the time of plate replacement and used to determine the average egg-laying time. When oviposition was evaluated in mated females, virgin females were collected following the indicated procedure, mated at day 1 and transferred to egg laying plates on day 4-5.

For hatchability assessment, plates with eggs were kept at 25 °C, and L1 larvae were counted daily until no newly hatched larvae were observed. Hatchability was expressed as the percentage of eggs that successfully completed embryonic development and hatched as larvae.

### Follicle classification

Ovaries were dissected in cold PBS, fixed for 30 min in 4% paraformaldehyde, rinsed with PBS containing Triton (0.3%) (PBST), stained with DAPI for 30 min, rinsed with PBS, and mounted in 50% PBS glycerol or Fluoromount-G (Southern Biotech). Follicles were staged following published criteria (Beard et al., 2023; Deady et al., 2017; Jia et al., 2016). Briefly, mature follicles were identified by their long dorsal appendage and by DAPI staining, as they lack nurse cell nuclei in the anterior region of the follicle. We classified stage 14 follicles as ‘early’ if they lacked nurse cells and their nuclei were randomly distributed, and as ‘fully mature’ if the nuclei were aligned in a row, as shown in **Fig. S16**. *dilp8-eGFP^MI00727^*is only active in fully mature stage 14 follicles (**Fig. 1**).

### Egg-laying time assay

The egg-laying process can be divided into three stages: ovulation, transit through the lateral and common oviduct, and transit through the uterus and oviposition. The total time required to lay a single egg can be estimated from the number of eggs laid per female per day. The duration of each stage can be inferred based on the assumption that the probability of finding an egg in a specific region of the reproductive tract at any given time is proportional to the duration of that stage. The time required for each stage is then calculated by multiplying the total egg-laying time by the frequency of eggs found within each reproductive tract region (Beard et al., 2023). To quantify the distribution of eggs within the reproductive tract, 5-d old virgin females were treated as in the Egg-Laying Assay, mated, and frozen at -80 °C for five minutes six hours after mating. They were then stored frozen until further analysis. Females were dissected, and the position of the egg within the reproductive tract was recorded as follows:

- None (N) – No egg found in the reproductive tract
- Bilateral (B) – More than 75% of an egg in the lateral oviduct
- Common (C) – Egg entirely within the common oviduct
- Uterus (U) – Egg located in the uterus
- Ejected (E) – Egg expelled from the uterus

To calculate ovulation time, oviduct time, and uterus time, the total egg-laying time was multiplied by the percentage of eggs in ovulation (%N), oviduct (%B + %C), and uterus (%U + %E), respectively. The egg-laying time is calculated using the number of eggs/female/day (Beard et al., 2023).

### Induction of egg retention

Egg retention was induced by placing females on hard-agar (5% agar) egg-laying plates without yeast paste. Briefly, virgin females were collected as described above and placed in vials with yeast-paste-supplemented food to induce oocyte production for four days. On day four, females were transferred to hard-agar egg-laying plates for seven days. Under these conditions, females laid very few eggs, regardless of genotype. On day 11 after eclosion, virgin females were mated and placed on standard egg-laying plates supplemented with yeast paste. Hatchability of laid eggs was evaluated on the first and third day after retention. Eggs laid during the first 24-h period corresponded to those retained in the ovary during the experimental retention period. Eggs laid on the third day were expected to be mostly produced after mating and, therefore, considered non-retained. In parallel, virgin females were collected in the same way but mated on day 1 after eclosion and continuously housed with males until the end of the experiment served as the “no-retention” control. They were kept on yeast-supplemented standard food for four days and transferred to regular egg-laying plates supplemented with yeast on day five. On day 11, they followed the same regime as retained females.

### Blockage of vesicle release in follicle cells

Virgin *dilp8-1-GAL4 UAS-shi^ts^ UAS-sGFP* females, were developed at room temperature (RT) and maintained at the same temperature after eclosion. On day 7, they were shifted to 30 °C for 24 h, or kept at RT as controls. Control flies without *UAS-shi^ts^*received the same temperature regime. Ovaries were dissected as indicated below and stained with a-Dilp8 and a-GFP antibodies.

To assess the effect of the blockage of vesicle release on egg laying, flies were developed at 18 °C. Virgin females were maintained at 30 °C on vials with regular food. On day five after eclosion, flies were transferred to egg-laying plates supplemented with a smear of yeast paste and allowed to lay eggs for 48 h. Egg-laying plates were replaced every 24 h at 30 °C, depending on the corresponding condition. Flies were assayed in groups of five individuals.

### Anti-DVGlut Antibody Production

A polyclonal antibody against DVGlut was generated using a recombinant C-terminal fragment (amino acids 561-632) of the protein as antigen as described in (Mahr and Abele, 2006). The selected immunogenic region corresponds to the C-terminal sequence:

GSEYTEQSQMQQSTAISYGATGHVANNPFAMASGAPPIAEEDAPPTYGDVTNPGQYGYTQGQMPSYDPQGYQQQ

The coding sequence for this DVGlut C-terminal fragment was cloned in-frame with glutathione S-transferase (GST) [pGEX4T1-GST DVGlut C-Terminal a gift from Hermann Aberle (Mahr and Abele, 2006)] to produce a soluble GST–DVGlut-C-terminal fusion protein. The fusion protein was expressed in *E. coli* and purified by glutathione affinity chromatography (SICGEN). The purified GST-fusion protein was used to immunize rabbits following a standard multi-boost polyclonal immunization protocol. Serum collected after the final boost was screened by ELISA and Western blot for reactivity against the immunizing fragment and Drosophila tissue extracts (Eurogentec). For increased specificity, the resulting antiserum was affinity-purified using an immobilized MBP-DVGlut-C-terminal fusion protein, containing the same C-terminal fragment fused to maltose-binding protein (MBP). The MBP-fusion protein was expressed and purified under native conditions and covalently coupled to an affinity matrix. Crude serum was passed through the immobilized antigen column, and DVGlut-specific antibodies were selectively eluted using low-pH glycine buffer, immediately neutralized, and dialyzed against PBS. This step ensured enrichment of high-specificity antibodies that recognize epitopes within the DVGlut C-terminal domain (SICGEN). The final affinity-purified antibody preparation showed strong immunoreactivity and specificity for DVGlut in ELISA, Western blot, and immunohistochemistry (**Fig. S12**).

### Immunofluorescence analyses

For **Fig. 1 and 4-6**, and **Fig. S3, S8, and S9**, ovaries were dissected in Schneider Medium (Biowest – cat. #L0207 or Gibco - cat. #21720-024), fixed for 30 min in 4% paraformaldehyde, rinsed with PBS with Triton (0.3%) (PBST), incubated with primary antibody for 24 h and with fluorescently labeled secondary antibody for 24 h in PBST with 1% bovine serum albumin. Samples were washed 3× for 30 min each in PBST after each antibody incubation. Nuclei were counterstained with DAPI (Sigma) and tissues were mounted in Fluoromount-G (Southern Biotech) or Dabco mounting medium (Sigma-Aldrich). Primary antibodies used were: Rat anti-Dilp8, 1:500 (Colombani, Andersen, and Léopold 2012), Rabbit Anti-GFP, 1:200 (Life Technologies, A11122), Mouse anti-GFP, 1:200 (Developmental Studies Hybridoma Bank (DSHB), DSHB-GFP-4C9), Rabbit Living Colors DsRed, 1:500 (Clontech, 632496), Rabbit anti-Tyrosine decarboxylase 2 (anti-Tdc2) 1:200 (Covalab, pab0822-P), and Rabbit anti-VGlut, 1:1000 (generated in this study). Secondary antibodies: Donkey Anti-Rat Alexa Fluor 647, 1:250 (Jackson 712-605-153), Goat Anti-Rabbit Alexa Fluor 488, 1:250 (Invitrogen A11008), Goat Anti-Mouse Alexa Fluor 568, 1:250 (Life Technologies, A11036), Donkey anti-rabbit Alexa Fluor 647, 1:250 (Jackson Immuno Research, 711-605-152); Goat Anti-Horseradish Peroxidase (HRP) Alexa Fluor 647, 1:250 (Jackson Immuno Research, 123-605-021). Images were acquired on a Leica SP5 laser scanning confocal microscope using a 10x or 20x dry objective. Ovaries from both virgin and mated females were prepared separately and we observed no obvious differences in the expression patterns tested.

For **Fig. S6 and S10**, adult brains, VNCs, and reproductive systems of 3 to 8 days old flies were dissected in cold phosphate-buffered saline (PBS) and immediately transferred to cold PFA 4% in PBL (PBS with 0.12 M Lysine) and fixed for 30 min at RT, washed three times for 5 min in PBT (PBS with 0.5% Triton X-100) and blocked for 30 min at RT in 10% normal goat serum in PBT (Sigma, cat# G9023). Samples were incubated with the primary antibodies in blocking solution, for 72 h at 4 °C. In adult brain and VNCs, the primary antibody Rabbit anti-GFP 1:1000 (Molecular Probes, A11122) was used together with Mouse anti-nc82 1:10 (DSHB, AB_2314866). Samples were washed three times for 5 min in PBT and incubated in Goat Anti-Rabbit Alexa Fluor 488 (Molecular Probes, A11034) and Goat Anti-Mouse Alexa Fluor 594 (Molecular Probes, A11032) secondary antibodies 1:500 for 72 h at 4°C. In reproductive systems, the same protocol was used with the same primary rabbit anti-GFP antibody followed by the secondary Goat anti-rabbit antibody together with Alexa 594-conjugated phalloidin (Molecular Probes, A12381). After incubation with secondary antibodies, all samples were washed three times for 5 min in PBT and mounted in VectaShield medium (Vector Laboratories, cat#H-1000). Images were acquired on a Zeiss LSM 980 laser scanning confocal microscope using a 25X immersion objective (Zeiss). After acquisition, colour levels were adjusted using Fiji for optimal display.

### qRT-PCR

The material used for the qRT–PCR experiments were obtained from 20 ovaries per biological replica from 5-day-old female virgins (Fig. 1C and Fig. S5), or from 5 whole virgin females or males aged 5 days per biological replica (Fig. 5H-I). The samples were macerated using pellet pestles and homogenized in 1 ml of NZYol (NZYTech) and centrifuged at 12,000 × g for 1 min, to lower tissue debris. After the centrifugation, half volume of absolute ethanol was added to the supernatant and mixed well. Then, the sample was loaded in a binding column of the NZY Total RNA Isolation kit (NZYTech). An extra DNAse treatment (Turbo DNA-free kit, Ambion, Life Technologies) was performed to reduce gDNA contamination. cDNA synthesis was performed using the Maxima First Strand cDNA Synthesis Kit for RT–quantitative PCR (Thermo Scientific) or NZY First-Strand cDNA Synthesis Kit, following manufacturer’s instructions. qRT–PCR experiments were performed as described previously (Heredia et al. 2021). Briefly, the experiments were performed in a LightCycler® 96 Instrument (Roche) or in a CFX96^TM^ (Bio-Rad) using the FastStart Essential DNA Green Master dye and polymerase (Roche) or the NZYSupreme qPCR Green Master Mix (2x) (NZYTech), respectively, depending on the experiment. The final volume for each reaction was 10 μl, consisting of 5 μl of dye and polymerase (master mix), 2 μl of cDNA sample (diluted to an estimated 1-10 ng/μl equivalent of RNA) and 3 μl of the specific primer pairs. The efficiency of the primers for the qPCR was verified by analyzing a standard curve with 3 serial dilutions of gDNA from w1118 animals and the production of primer dimer was checked by analyzing the melting curve. Only primers with more than 90% of efficiency and with no primer dimer formation were used. qRT-PCR results were expressed as mRNA levels relative to *rp49* (*rp49* = 1). Primer pairs used for *D. melanogaster* genes:

*rp49*_F 5′TTGAGAACGCAGGCGACCGT 3′

*rp49*_R 5′CGTCTCCTCCAAGAAGCGCAAG 3′

*dilp8*_F 5′CGACAGAAGGTCCATCGAGT 3′

*dilp8*_R 5′GATGCTTGTTGTGCGTTTTG 3′

*dilp3*_F 5′CCGAAACTCTCTCCAAGCTC 3′

*dilp3*_R 5′ GCCATCGATCTGATTGAAGTT 3′

*CG15279*_F 5’ TCGTCTATCCAAATTGGTCTTACTC 3’

*CG15279*_R 5’ AATGGCCACGATCATCCA 3’

*CG1698*_F 5’ GTCTACGACTTTTCACCCATCA 3’

*CG1698*_R 5’ AACGTGGCGTAGTAGGTGGT 3’

*CG3759*_F 5’ AAACTGTAGGCGAATCAGCAG 3’

*CG3759*_R 5’ CTCCTTTCGATAAGTGGTGTTGT 3’

*CG6508*_F 5’ AGGTCCAGGCGGATAACAAT 3’

*CG6508*_R 5’ TTACCCTGCAAGTATGTGAAGC 3’

*Oamb*_F 5’ GGAGAAAACTGGCCAGAACTAA 3’

*Oamb*_R 5’ TTACCCTGCAAGTATGTGAAGC 3’

### Western Blot

For the experiments described in **Figs S14, S15**, female pupae reared at controlled conditions were transferred to a new vial and fed on food supplemented with yeast paste for four days after eclosion. On day four, flies were separated into two groups. Control flies were kept in vials with yeasted food while starved flies were transferred to 2% agar in water. One day later, for **Fig. S14C, S15 (bottom)**, the thorax and head were dissected in ice-cold PBS and stored in 250 μl of ice-cold modified RIPA buffer with NaF, NaVO4 and SpI proteases and phosphatases inhibitors. Each sample contained material from five female flies. Protein concentration was measured with Bradford assay and 30 μg of protein separated on a 12% SDS-PAGE gel, transferred and analyzed with a Rabbit monoclonal anti-Akt (pan) antibody 1:1000 (C67E7) (Cell Signaling, #4691) and Rabbit monoclonal anti-Phospho-Akt 1:1000 (Ser473) (D9E) (Ser405 in *Drosophila*; Yang et al., 2006) (Cell signaling, #4060). An Donkey anti-rabbit ECL Horseradish peroxidase (HRP) conjugated was used as secondary antibody 1:5000 (NA934, Amersham). P-Akt/Akt was calculated as the ratio between the both p-Akt bands, over both total Akt bands, which Akt forms generated by alternative splicing (Yang et al., 2006). In this experiment, Rabbit polyclonal anti-α-Actin-1 (ACTA1) antibody 1:2000 (A2066, Sigma) served as loading control. For **Fig. S14A, S15 (top)**, one day later, the thorax and head were dissected in ice-cold PBS and stored in 100 μl of ice-cold 2X SDS PAGE loading buffer. Each sample contains material from five female flies. Protein concentration was measured in NanoDropTM One and 75 μg of protein separated on a 10% SDS-PAGE gel, transferred, and analyzed as above. Mouse monoclonal anti-alpha-tubulin 1:1000 (DSHB, 12G10) served as loading control. For **Fig. S12A-B**, western blots were analyzed as described in the figure legend.

### Statistical analysis

For all tests, alpha was set at 0.05 a priori. For quantitative data, comparison between multiple conditions was done using ANOVA when samples had normal distribution (Shapiro-Wilk test and equal variance). If the result of these tests was statistically significant, then Tukey’s HSD post-hoc tests were applied for the assessment of statistical significance of pairwise comparisons. When samples had unequal standard deviations, the Brown-Forsythe and Welch ANOVA and Dunnett’s T3 test was used instead. Comparisons between two conditions were done using two-tailed unpaired (unless noted otherwise) Student’s t-test, when samples had normal distribution and equal variance. Comparison between average proportions was performed using Student’s t –test with Bonferroni correction. Z-score test was used for egg-laying time analysis.

## AUTHORS CONTRIBUTIONS

Y.V., F.H., R.Z., J.M., M.S.P., M.G., L.L., L.S.C., A.P.C., M.L.V, A.G., and A.M.G. performed genetic, phenotypic, molecular, and behavioral experiments. F.H., C.H., M.L.V, A.G., and A.M.G. designed research. C.H. generated and characterized the anti-DVGlut antibody. F.H., A.G., and A.M.G., supervised the work. Y.V., F.H., R.Z., C.H., M.L.V, A.G., and A.M.G. contributed to image and data analyses. A.G. and A.M.G. conceived the study and wrote the manuscript with the help of all authors.

## ACKNOWLEDGMENTS

We thank Élio Sucena for the Mel2 population, Pierre Léopold for the a-Dilp8 antibody, Rita Teodoro for the anti-HRP antibody and stocks, Carlos Ribeiro for stocks, Hermann Aberle for the pGEX4T1-GST DVGlut C-Terminal plasmid, José Ramalho from SICGEN for help with the antibody generation, Maria Dominguez for discussions and support in the initial stages of this work, and Ana Margarida Moreira and Maria João Cruz for help in initial experiments of this work. We thank lab members for discussions and the Jianjun Sun and team for discussions and for sharing unpublished results prior to publication. We thank the Champalimaud Foundation fly facility and imaging facility for support. Stocks obtained from the Bloomington Drosophila Stock Center (NIH P40OD018537), the Kyoto Drosophila Stock Center, and the Vienna Drosophila Resource Center (VDRC) at Vienna BioCenter Core Facilities (VBCF), member of the Vienna BioCenter (VBC), Austria, were used in this study. Work in the Integrative Biomedicine Laboratory was supported by the Fundação para Ciência e Tecnologia (FCT; PTDC/BIA-BID/31071/2017; EXPL/BIA-BID/1524/2021; EXPL/BIA-COM/1296/2021; 10.54499/2022.03859.PTDC; 2023.15344.PEX), by the Research Units iNOVA4Health (10.54499/UIDB/04462/2020) and cE3c - Centre for Ecology, Evolution and Environmental Changes (10.54499/UIDB/00329/2020), by the The Associate Laboratory Life Sciences for a Healthy and Sustainable Future (LS4FUTURE) and CHANGE financed by the FCT/MCTES (Ministério da Ciência, Tecnologia e Ensino Superior), by Congento LISBOA-01-0145-FEDER-022170, co-financed by FCT/Lisboa2020; UID/Multi/04462/2019; and by the Microscopy Facility of the Faculdade de Ciências da Universidade de Lisboa (PPBI-POCI-01-0145-FEDER-022122). Work in the Garelli lab was supported by Agencia Nacional de Promoción Científica y Tecnológica (ANPCyT; PICT 2017-0254 and PICT 2020-01568), Consejo Nacional de Investigaciones Científicas y Técnicas (CONICET; PIP11220150100182CO) and Universidad Nacional del Sur PGI 24/B288. CH was supported by the FCT (PTDC/BIA-BID/0681/2021-ZeroToNeuron). MLV was supported by the Champalimaud Foundation. AMG and FH were individually supported by grants 10.54499/CEECINST/00102/2018/CP1567/CT0031 and 10.54499/DL57/2016/CP1457/CT0016, respectively. AG and YV are CONICET researchers, MSP holds a CONICET PhD fellowship. All data are available in the manuscript or the supporting information. All reagents and fly strains generated in this study are available from the corresponding authors without restriction

