## Supplementary Material for "Dilp8 relaxin signaling from ovarian follicle cells to Lgr3+ neurons promotes spontaneous ovulation and oocyte quality in *Drosophila*"

c iNOVA4Health, Nova Medical School, NMS, Universidade Nova de Lisboa, Lisbon 1150-082, Portugal

d Champalimaud Research, Champalimaud Center for the Unknown, Lisbon 1400-038, Portugal.

1 These authors contributed equally to this work.

**The PDF file includes:**
Table S1 to S2
Figures S1 to S16
Supplemental References

| | **Virgin** | | **Mated** | |
|---|---|---|---|---|
| **Egg-laying time (min.)** | ***dilp8 +/+*** | ***dilp8 -/-*** | ***dilp8 +/+*** | ***dilp8 -/-*** |
| Ovulation | 66.11 ± 10.36 | 221.24 ± 105.73 *** | 21.66 ± 5.40 | 31.42 ± 13.06 (ns) |
| Oviduct | 13.84 ± 8.33 | 65.07 ± 57.13 (ns) | 4.56 ± 5.50 | 5.50 ± 4.35 (ns) |
| Uterus | 3.07 ± 4.19 | 13.01 ± 25.51 ** | 24.51 ± 29.06 | 29.06 ± 12.35 (ns) |
| **Total** | 83.02 | 299.32 | 50.73 | 65.98 |

**Table S1.** Ovulation time, Oviduct transit time and Uterus residency time were calculated as described in Methods. Asterisks indicate $P$ value: (***): $P < 0.0005$, (**): $P < 0.005$, Z-score test

| probeset_id | ID1 | ID2 | Stg10A | Stg10B | Stg12 | Stg14 |
|---|---|---|---|---|---|---|
| 1626000_at | aspartic peptidase | CG6508 | 1 | 0.86 | 1.00 | 24.20 |
| 1623083_at | CG1698 | CG1698 | 1 | 1.11 | 1.21 | 22.89 |
| 1625664_at | CG14059 | ilp8 | 1 | 0.83 | 1.00 | 17.41 |
| 1635328_at | CG3856 | Oamb | 1 | 1.07 | 1.01 | 16.94 |
| 1626831_at | atilla | CG6579 | 1 | 0.97 | 0.86 | 16.60 |
| 1636888_a_at | CG1462 | Aph-4 | 1 | 0.98 | 0.89 | 14.05 |
| 1627590_at | CG9336 | CG9336 | 1 | 0.92 | 1.17 | 10.43 |
| 1636311_at | CG9042 | Gpdh | 1 | 1.05 | 1.24 | 9.16 |
| 1637338_at | chorion | CG12517 | 1 | 0.94 | 1.16 | 8.40 |
| 1623536_s_at | CG11347 | CG11347 | 1 | 0.68 | 0.94 | 7.03 |
| 1636595_a_at | CG13138 | CG13138 | 1 | 0.99 | 0.94 | 6.44 |
| 1633268_s_at | CG10806 | Nha1 | 1 | 0.92 | 0.88 | 5.71 |
| 1624371_a_at | transglutaminase | CG7356 | 1 | 0.96 | 0.85 | 5.53 |
| 1638462_at | alkaline phosphatase like | CG18088 | 1 | 0.92 | 1.21 | 4.85 |
| 1636835_at | GABA simporter | CG16700 | 1 | 0.88 | 1.11 | 4.36 |
| 1632160_s_at | cation amino symporter | CG15279 | 1 | 1.10 | 1.11 | 4.26 |
| 1627493_at | CG5958 | CG5958 | 1 | 0.50 | 0.87 | 4.18 |
| 1632932_a_at | multiplexin | CG33171-RC | 1 | 1.08 | 1.03 | 3.41 |
| 1623299_at | CG1794 | Mmp2 | 1 | 0.91 | 1.22 | 3.31 |
| 1639733_s_at | CG14275 | CG14275 | 1 | 1.00 | 1.05 | 3.14 |
| 1631441_s_at | slowdown EGF like | CG7447 | 1 | 0.82 | 0.90 | 2.82 |
| 1640303_a_at | CG8588 | pst | 1 | 0.84 | 0.93 | 2.79 |
| 1624435_at | CG12143 | Tsp42Ej | 1 | 0.97 | 0.95 | 2.48 |
| 1634388_at | CG30197 | CG30197 | 1 | 0.65 | 1.01 | 2.47 |
| 1624169_at | CG13317 | Ilp7 | 1 | 1.05 | 0.97 | 2.47 |
| 1639480_at | CG6871 | Cat | 1 | 0.93 | 0.96 | 2.41 |
| 1634935_a_at | CG1648 | CG1648 | 1 | 0.95 | 0.92 | 2.37 |
| 1625178_at | CG31038 | CG31038 | 1 | 1.23 | 0.90 | 2.36 |
| 1629095_a_at | CG10120 | Men | 1 | 1.00 | 0.89 | 2.35 |
| 1635449_s_at | CG9285 | Dip-B | 1 | 1.00 | 0.88 | 2.28 |
| 1640979_at | CG1681 | CG1681 | 1 | 0.92 | 1.00 | 2.14 |
| 1638556_s_at | CG3856 | Oamb | 1 | 0.93 | 1.07 | 2.14 |
| 1627744_at | CG15209 | CG15209 | 1 | 1.13 | 1.05 | 2.03 |
| 1631830_at | HDC03331 | NA | 1 | 0.96 | 1.20 | 2.02 |
| 1625245_at | CG3759 | Multicopper oxidase-1 | 1 | 0.99 | 0.99 | 1.95 |

**Table S2.** List of genes expressed in follicles that specifically increase their transcription in stage 14 follicles. Data were obtained from Tootle et al. (2011) and normalized to expression in stage 10A follicles. The table lists only those genes that show at least a two-fold increase in expression.

**Supplementary Figure S1.**

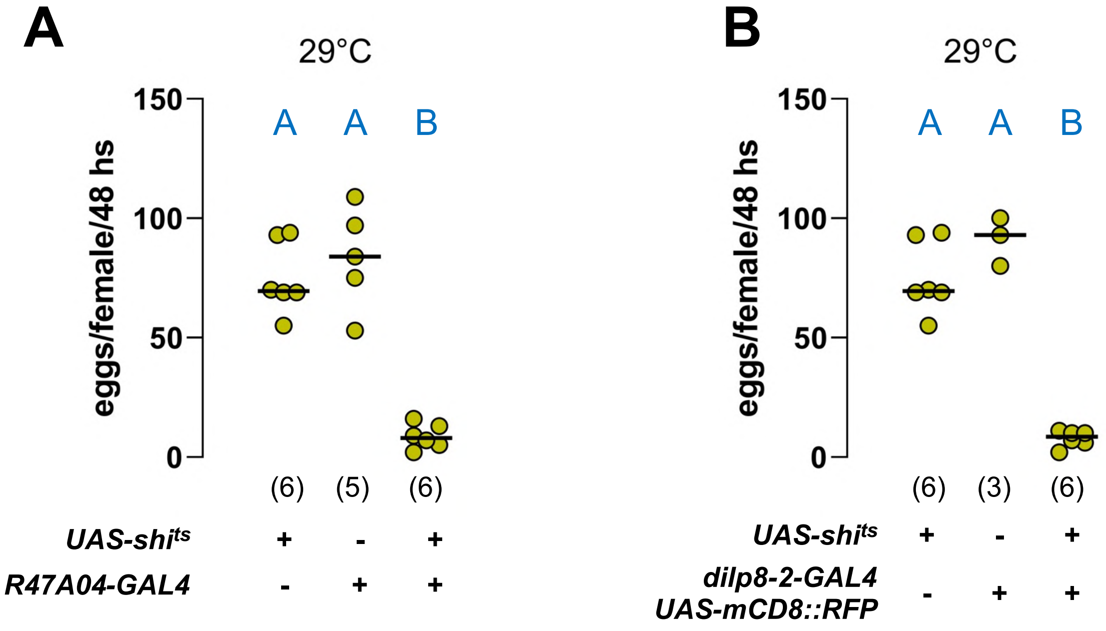


**Fig S1. Blocking the secretory pathway in stage 14 follicle cells decreases virgin egg laying. (A)** Expression of *UAS-shi^ts* in follicle cells with *R47A04-GAL4* or **(B)** *dilp8-2-GAL4* decreases the number of unfertilized eggs laid by virgin females at the restrictive temperature. Each dot represents an independent biological replicate (*N* in brackets) with five flies each. Therefore, the total number of flies is *N**5 for each condition. Horizontal line, median value. Same blue letter, $P > 0.05$, One-way ANOVA and Tukey's HSD test. The *UAS-shi^ts* data is the same for both panels.

**Supplementary Figure S2.**

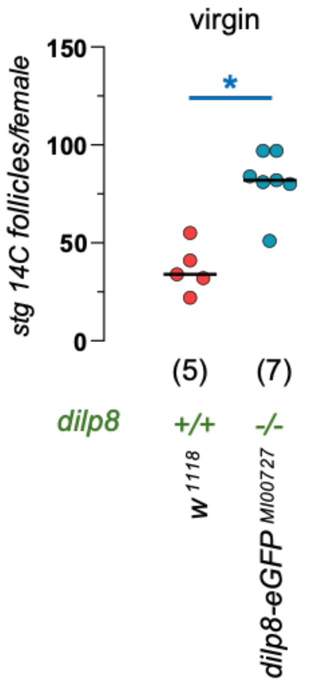


**Fig S2. *dilp8-eGFP$^{MI00727}$* mutants retain more mature follicles (stage 14 follicles) in the ovary.** The number of mature oocytes (stage 14 follicles) per adult (5- days old) female virgin was counted in *w$^{1118}$;; dilp8-eGFP$^{MI00727}$* homozygous mutants and *w$^{1118}$* controls. *dilp8-eGFP$^{MI00727}$* mutants retain significantly more stage 14 follicles in their ovaries than their controls. Each dot represents a fly. Horizontal line, median value. *, *P* = 0.0003, unpaired two-tailed Student's t-test.

**Supplementary Figure S3.**

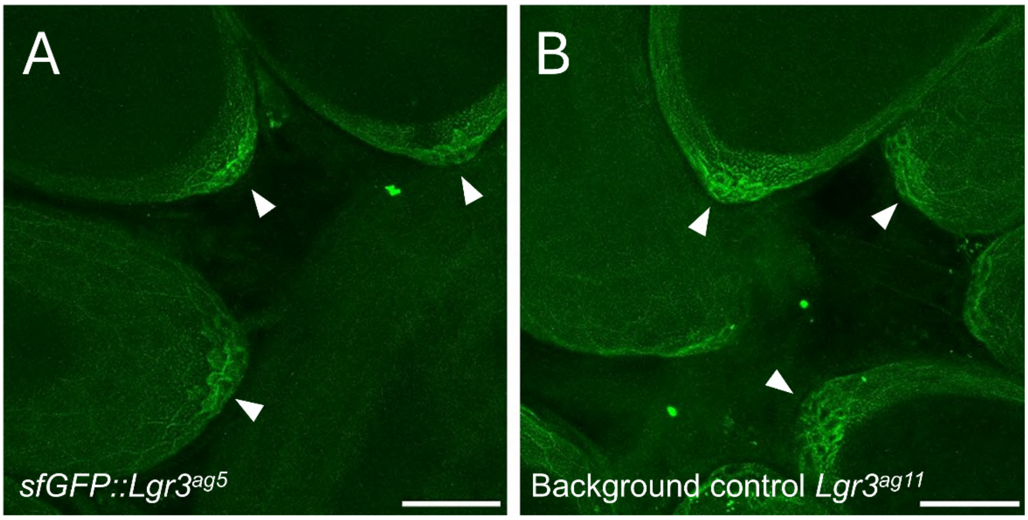


**Fig. S3. Posterior follicle cells (adjacent to the future aeropyle) do not express detectable Lgr3 protein. (A)** The fluorescent signal observed at the posterior tip of stage 14 follicles in flies carrying a translational reporter for Lgr3 expression (*sfGFP::Lgr3$^{ag5}$*) is not specific, as evidenced by the same signal appearing in **(B)** background control samples from flies lacking any GFP insertion (*Lgr3$^{ag11}$*). We conclude that the signal arises due to autofluorescence of the aeropyle, a chorionic structure. Scale bar = 100 µm.

**Supplementary Figure S4.**

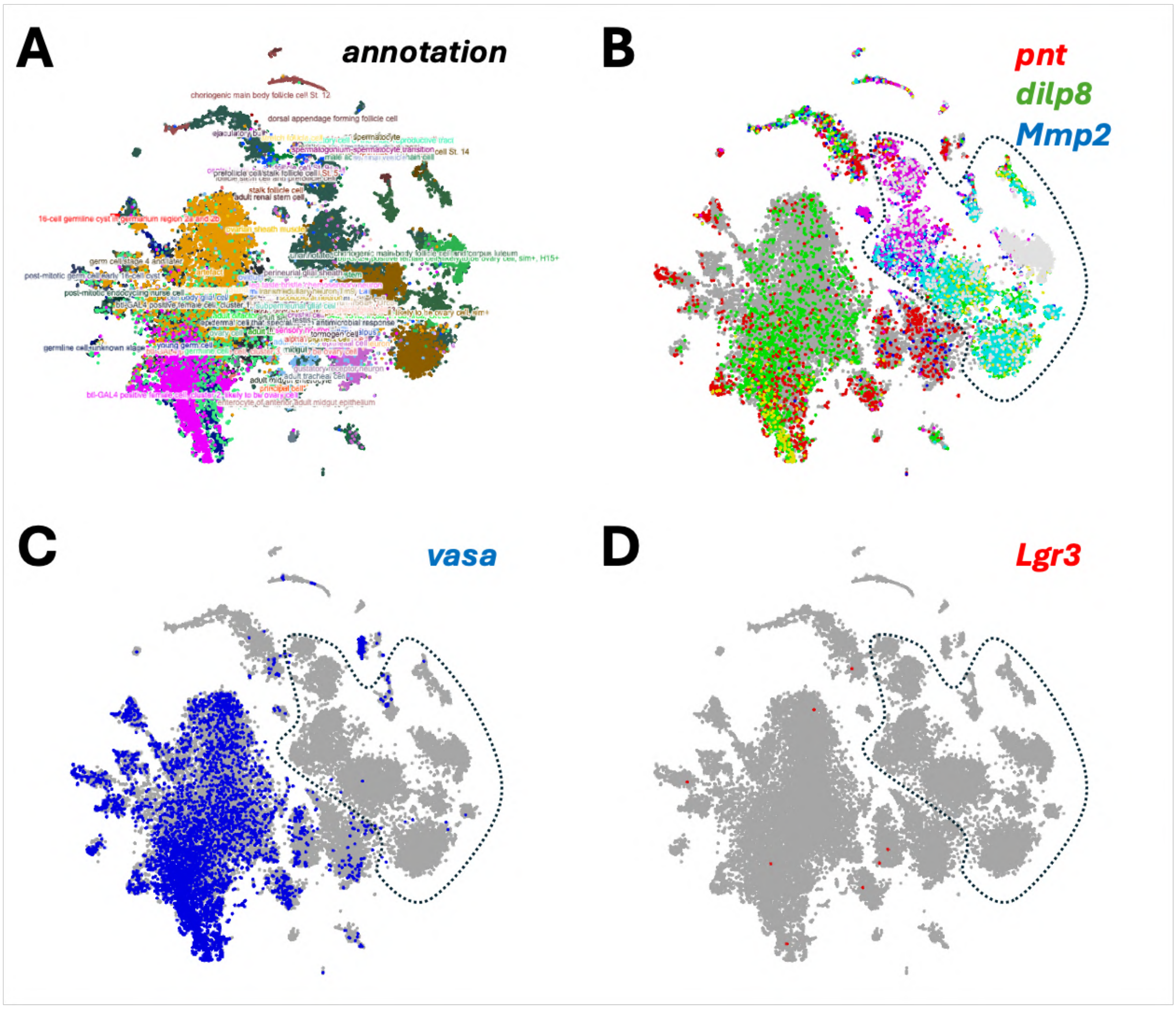


**Fig. S4. 10x-cross tissue single-cell RNA sequencing analyses of *Lgr3* expression in the female reproductive tract. (A)** Scope annotations of possible cell types. **(B)** *dilp8* (green), *pnt* (red), and *Mmp2* (blue) are used as markers of late-stage follicle cells. Cells triply positive for these markers are labeled in light gray and are found in clusters surrounded by the dotted line. **(C)** The germline marker, vasa (blue), is used to mark putative germline cells. **(D)** *Lgr3* expression (red), is very low in the female reproductive tract and is not detectable in late-stage follicle cells. Scope analyses (https://scope.aertslab.org/) of 10x-crosstissue FlyCellAtlas data (Li et al., 2022), female reproductive tract data, stringent. All gene expression levels were saturated, so even cells with minimal counts per million are labelled in these graphs.

**Supplementary Figure S5.**

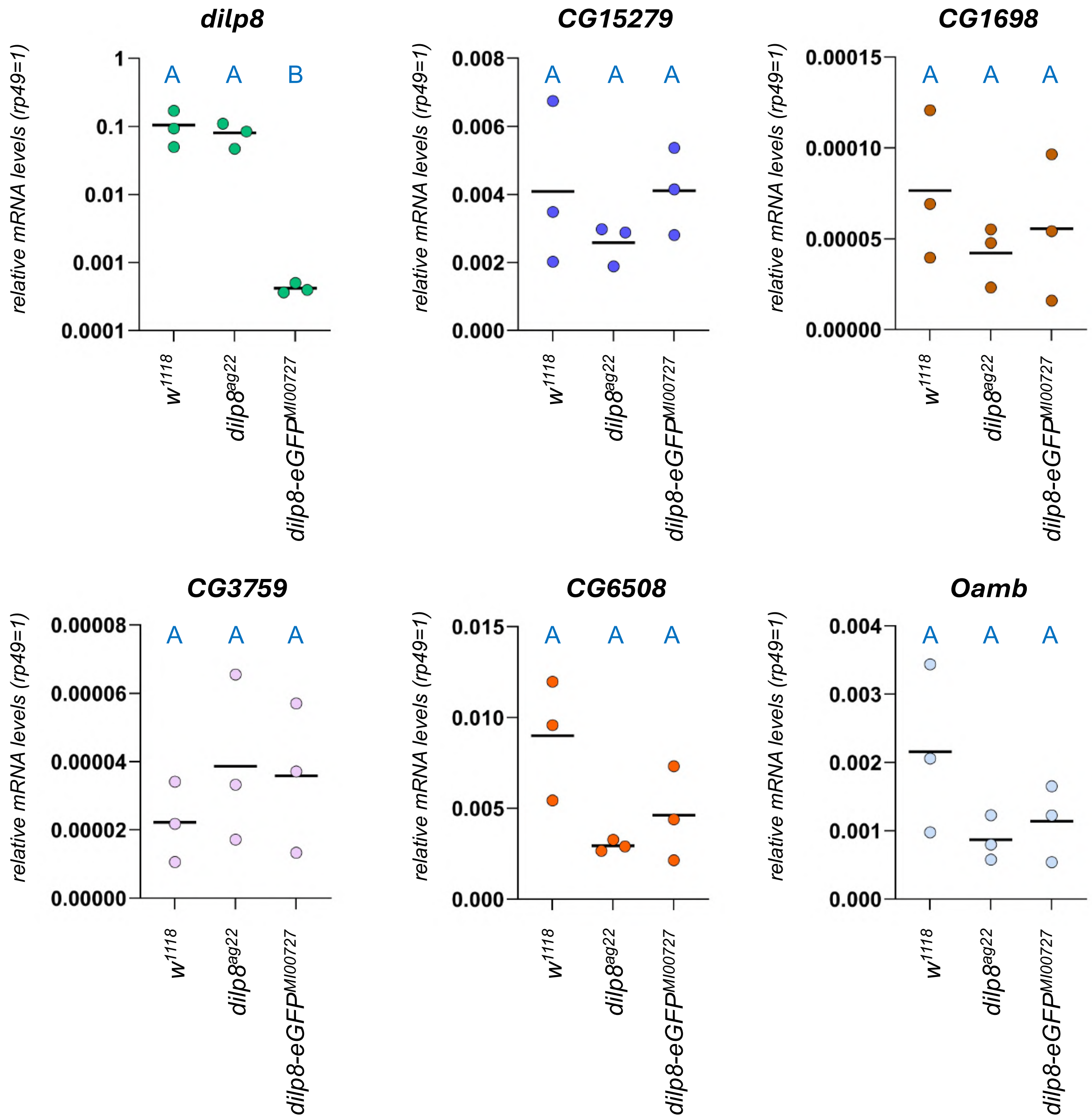


**Fig. S5. Expression of stage 14 follicle specific genes is not consistently affected in *dilp8* mutant.** Expression of a set of stage 14 follicle specific genes was evaluated by qRT-PCR in control ($w^{1118}$ and $dilp8^{ag22}$) and $dilp8\text{-}eGFP^{MI00727}$ mutant flies. Each dot represents an independent biological replicate ($N$ = 3) with three technical replicates each. Same blue letter, $P > 0.05$, ANOVA and Tukey's HSD test. The *dilp8* data is the same as that shown in Figure 1C, and is reproduced here for comparison.

**Supplementary Figure S6.**

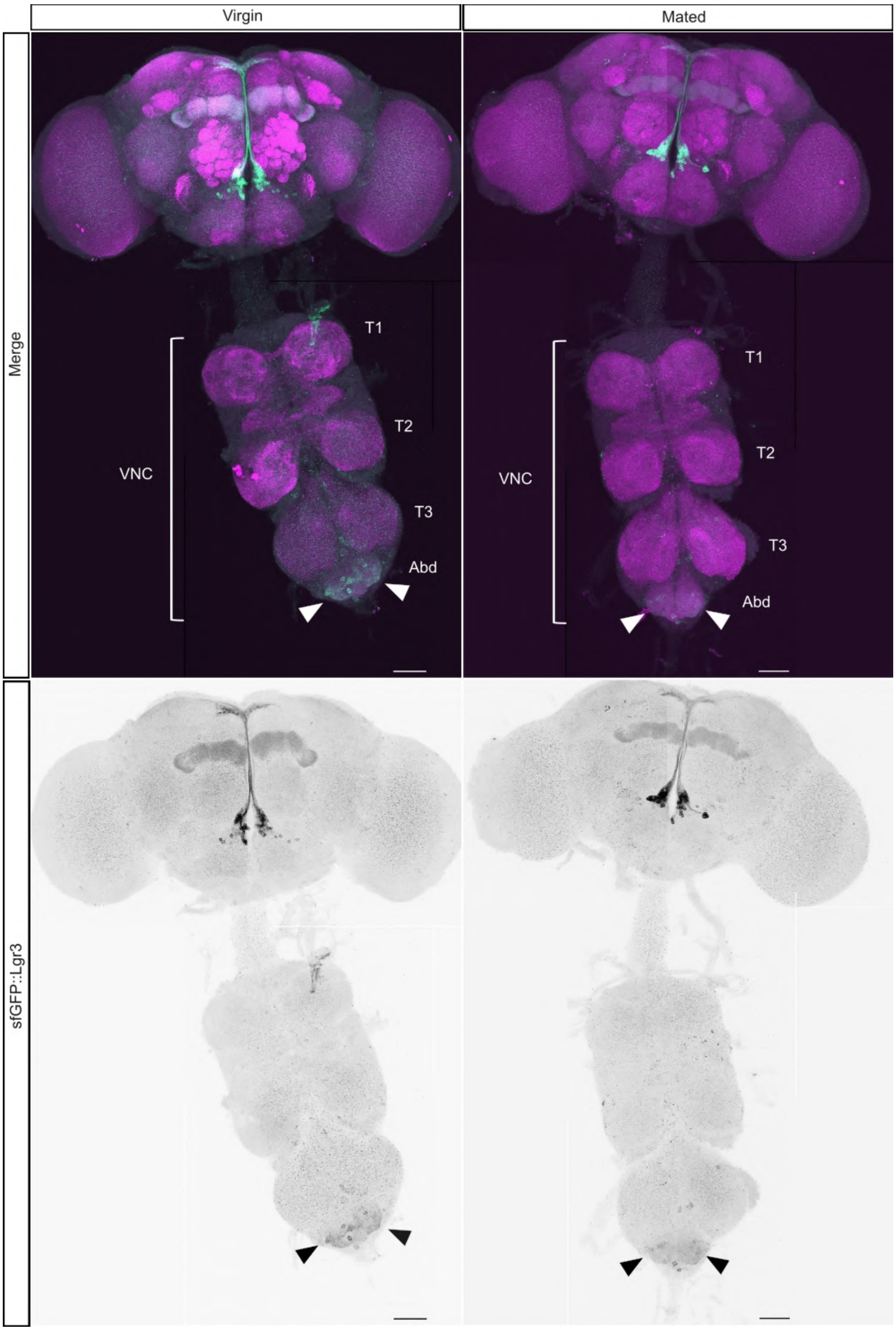


**Fig. S6. sfGFP::Lgr3 expression in the adult central nervous system of virgin and mated females.** Depicted are stitched assemblies of maximum intensity projections of confocal stacks obtained from central nervous system immunohistochemistry preparations of **(A)** virgin and **(B)** mated females expressing sfGFP::Lgr3, detected with an anti-GFP antibody (green (top) and black, single channel, bottom panels). In the top panels, the neuropil is counterstained with an anti-Bruchpilot antibody (anti-nc82, magenta). Thoracic segments T1-3 and the abdominal (Abd) ganglion are depicted. Arrowheads depict the region with sfGFP::Lgr3-positive cells in the Abd. The sfGFP::Lgr3-positive cells in the brain are similar to the sexually-dimorphic median bundle Lgr3-GAL4-positive cells described in Meissner et al. (2016). Scale bar = 50 μm.

**Supplementary Figure S7.**

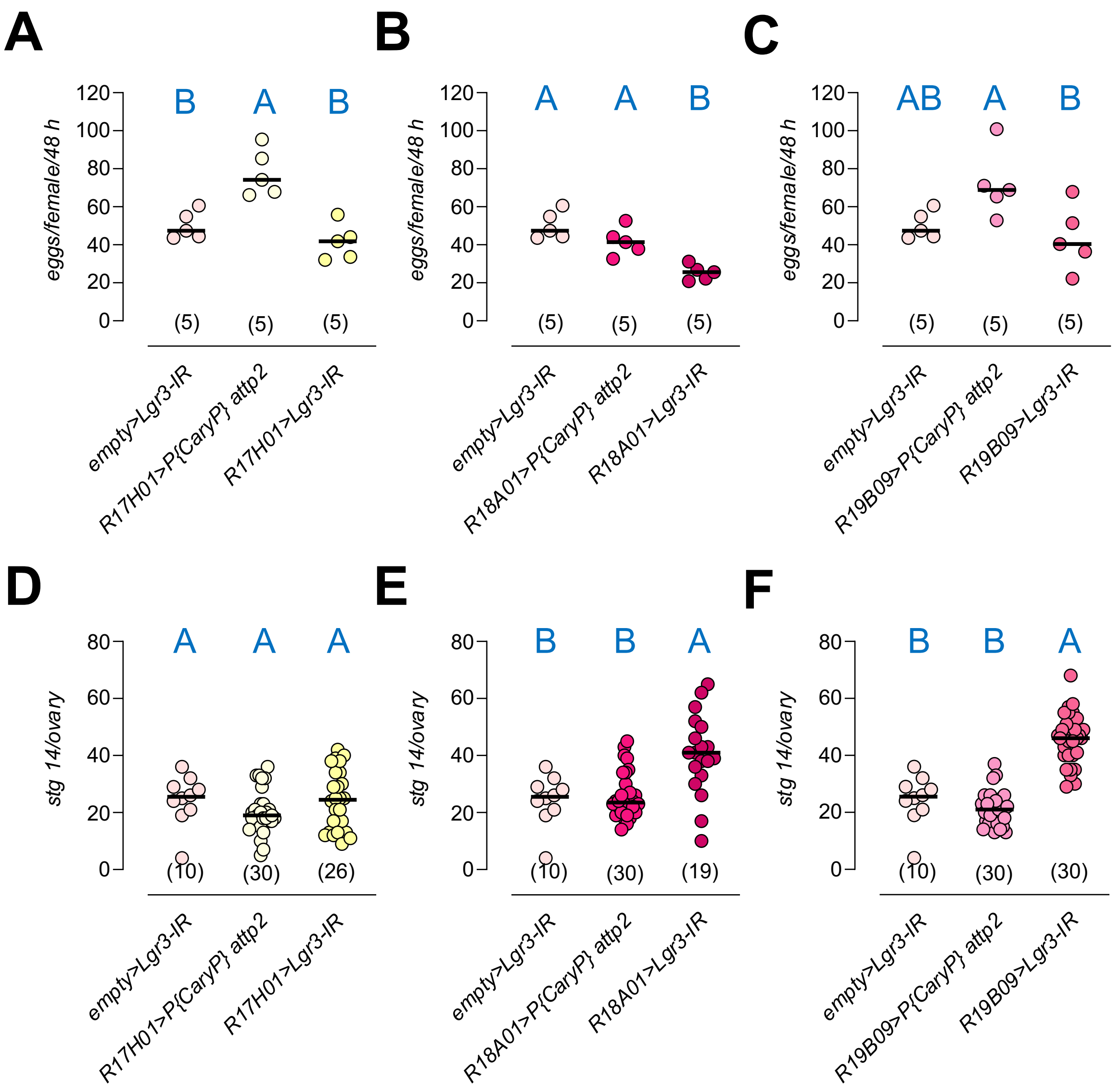


**Fig. S7. GAL4-alone controls for *R17H01*, *R18A01*, and *R19B09* drivers in Fig. 4D-E. (A-C)** Quantification of number of unfertilized eggs laid in animals expressing *Lgr3* knockdown in *R17H01>*, *R18A01>*, or *R19B09>* cells. **(D-F)** Same as (A-C), but for mature stage 14 follicles per ovary (stg 14/ovary). Data for *empty>Lgr3-IR* and *driver>Lgr3-IR* are the same as in Fig. 4D-E and are reproduced here for comparison. Each dot represents an independent biological replicate (*N* in brackets) with five flies per replicate. Horizontal line, median value. Same blue letter, $P > 0.05$, One-way ANOVA and Tukey's HSD test.

**Supplementary Figure S8.**

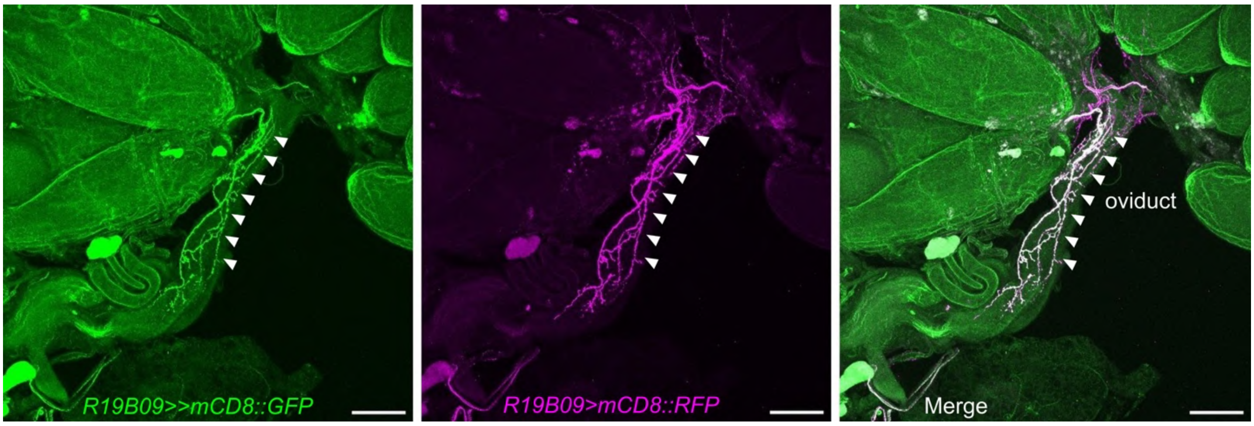


**Fig. S8. *R19B09-LexA* (*R19B09>>*) and *R19B09-GAL4* (*R19B09>*) drive expression in the same neurons that innervate the female reproductive tract.** Depicted are maximum intensity projections of confocal stacks of an immunohistochemistry preparation of a dissected reproductive system from a mated female. *R19B09-GAL4* (*R19B09>*) driving *UAS-mCD8::GFP* (*R19B09>mCD8::GFP*, green) and *R19B09-LexA* (*R19B09>>*) driving *LexAop2-mCD8::RFP* (*R19B09>>mCD8::RFP*, magenta) were detected with antibodies against GFP and RFP, respectively. Merge (right panel). Arrowheads depict the neurons innervating the common oviduct which are more evident in this image. Scale bar = 100 μm.

**Supplementary Figure S9.**

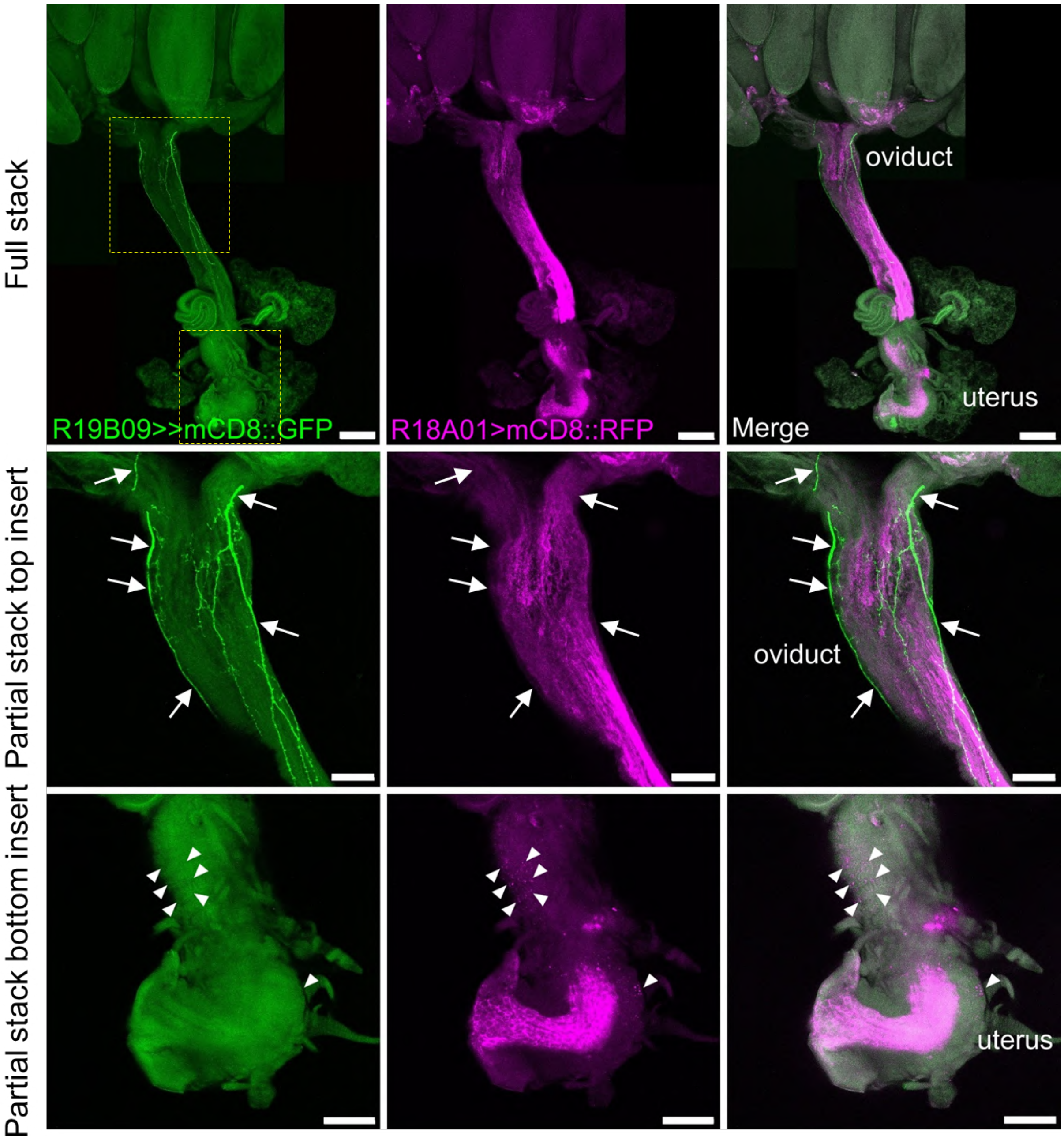


**Fig. S9. *R19B09-LexA* (*R19B09>>*) and *R18A01-GAL4* (*R18A01>*) do not drive expression in the same neurons that innervate the female reproductive tract.** Depicted are maximum intensity projections of confocal stacks of an immunohistochemistry preparation of a dissected reproductive system from a mated female. *R18A01-GAL4* (*R18A01>*) driving *UAS-mCD8::RFP* (*R18A01>mCD8::RFP*, magenta) and *R19B09-LexA* (*R19B09>>*) driving *LexAop2-mCD8::GFP* (*R19B09>>mCD8::GFP*, green) were detected with antibodies against GFP and RFP, respectively. Merge (right panel). Arrows depict *R19B09>>*-positive oviduct projections, which are negative for *R18A01>*-driven expression. Arrowheads depict the neuromuscular junctions on the uterus muscular wall that are positive for *R18A01>* and negative for *R19B09>>.* The neuromuscular junctions are very faint, because the *R18A01>* driver here is used in as a single copy. When two copies are used, the neuromuscular junction pattern and overall neuronal expression are much stronger and evident (see, e.g., **Fig. 6B”**). Scale bars = 100 µm, top panels; 50 µm, bottom inserts.

**Supplementary Figure S10.**

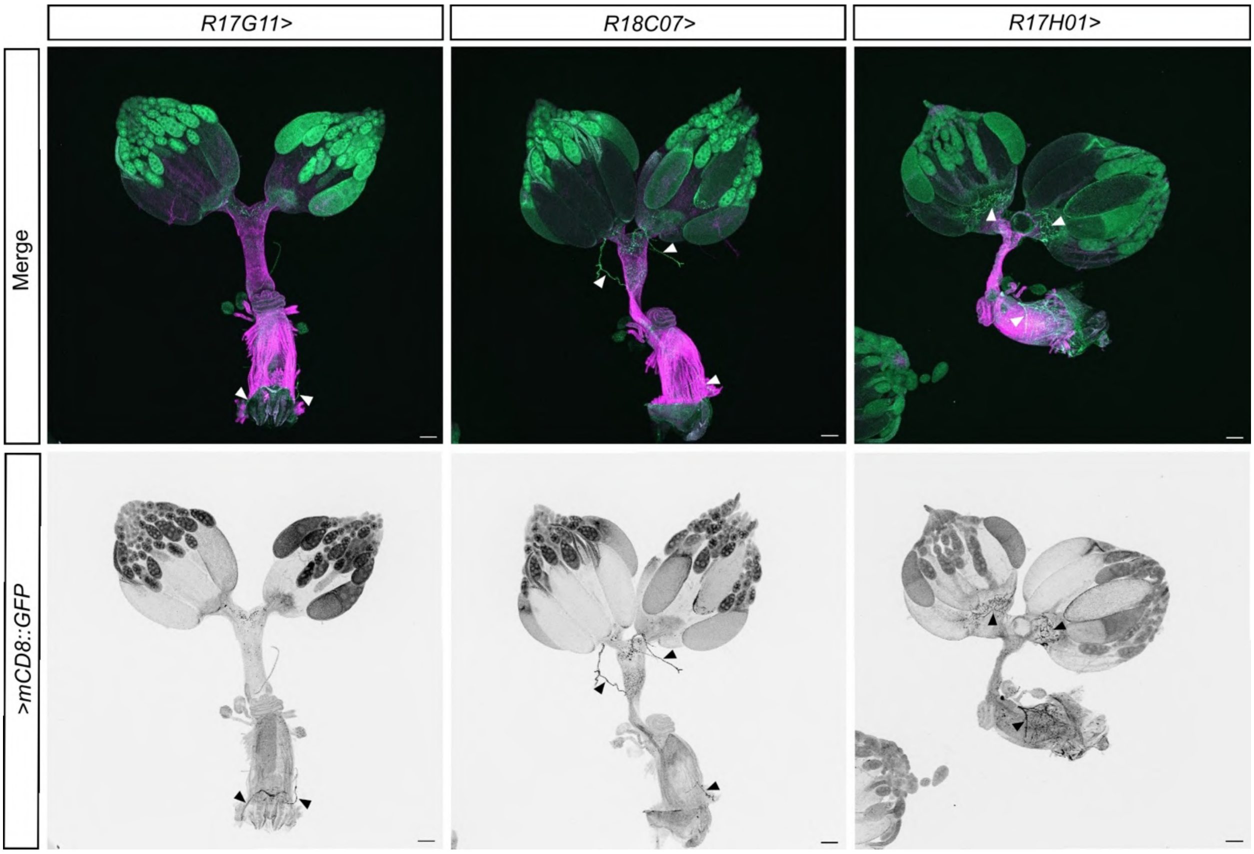


**Fig. S10. *Expression pattern of the Lgr3 cis-regulatory modules R17G11>*, *R18C07>*, and *R17H01>* in the female reproductive tract.** Depicted are maximum intensity projections of confocal stacks of an immunohistochemistry preparation of a dissected reproductive system from animals of the depicted genotypes driving *>mCD8::GFP* (green), and counterstained with phalloidin (magenta). Arrowheads depict possible neuronal fibers. Scale bar = 100 µm.

**Supplementary Figure S11.**

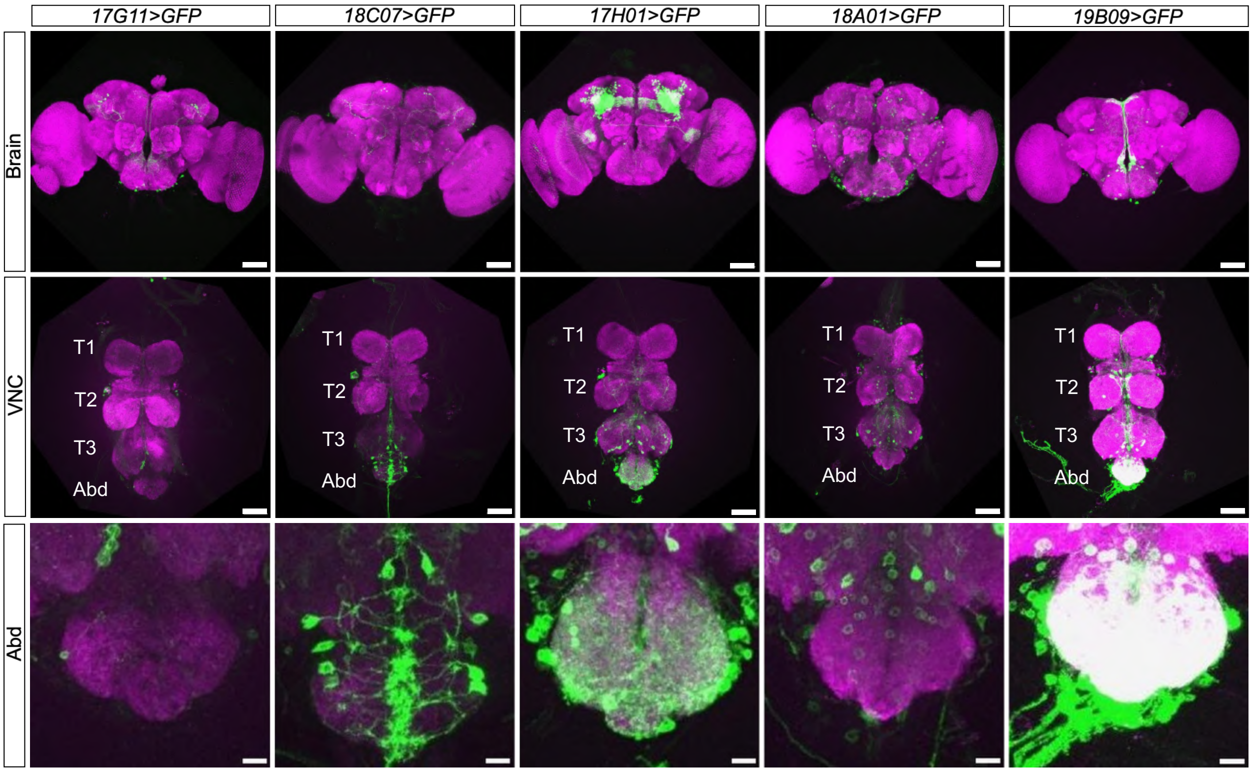


**Fig S11. FlyLight expression pattern of *Lgr3 cis-regulatory module GAL4* lines in the Ventral Nerve Cord of adult *D. melanogaster.*** All depicted *Lgr3 cis-regulatory module GAL4* lines drive at least some detectable expression in the adult brain (top roow), VNC (thoracic segments labeled as T1-3; middle row), and likely also in neurons of the abdominal ganglia (Abd; bottom row) (*UAS-mCD8::GFP*, green). The neuropile is marked in magenta. Both *R18A01-GAL4* and R19B09-GAL4, which drive expression in a subset of neurons where *Lgr3* is required for virgin unfertilized egg-laying and/or mature oocyte retention, are expressed in the Abd, which harbors neurons known to innervate the female reproductive tract. Scale bars, top and middle rows = 100 µm; bottom row = 20 µm. Images from https://flweb.janelia.org/ (Jenett et al., 2012; Pfeiffer et al., 2008).

**Supplementary Figure S12.**

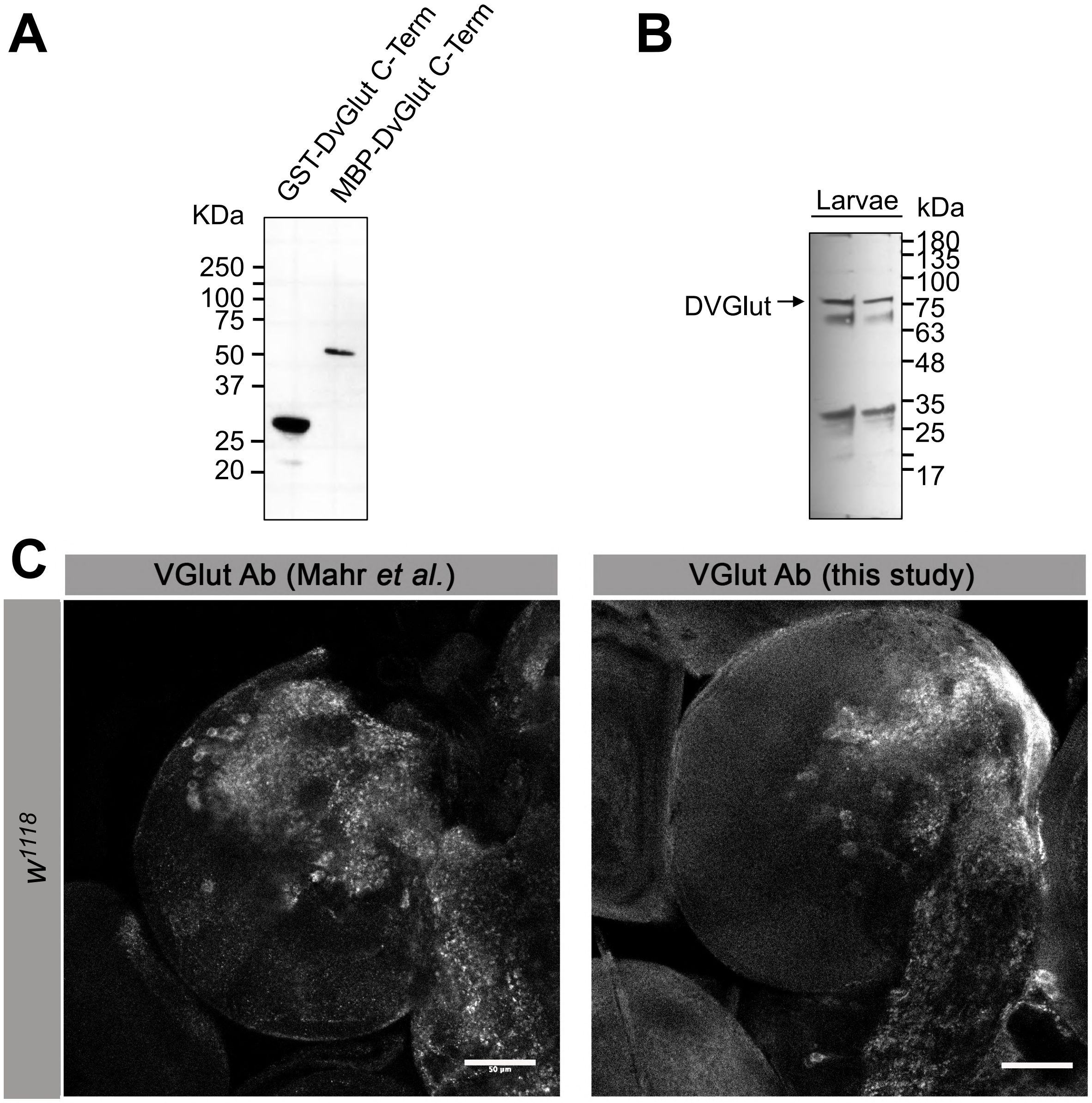


**Fig S12. Characterization of an Anti-DVGlut antibody generated in this study based on the same Drosophila VGlut1 epitope described in Mahr and Abele (2006). (A)** Western blot using Anti-DVGlut antibody at 1/1,000 dilution; lanes were loaded with 20-30 ng of recombinant fusion proteins. The secondary antibody used was goat polyclonal to rabbit IgG (DyLight®633) at 1/5,000 dilution. **(B)** Western blot showing that the Anti-DVGlut antibody detects an endogenous band migrating at the predicted molecular weight of *Drosophila* VGlut1.Anti-DvGlut Ab was used at a at 1/1,000 dilution; lanes were loaded with 10-20 µg of Drosophila brain extracts. Secondary antibody used was a goat polyclonal to rabbit IgG (DyLight®633) at 1/5,000 dilution. **(C)** Single confocal slice of a dissected central nervous system (brain region) from a $w^{1118}$ larva stained with anti-DVGlut (right) shows the expected pattern of glutamatergic neuron projections in the neuropil which are very similar to the image published by Mahr and Abele (2006). Scale bar 50 µm.

**Supplementary Figure S13.**

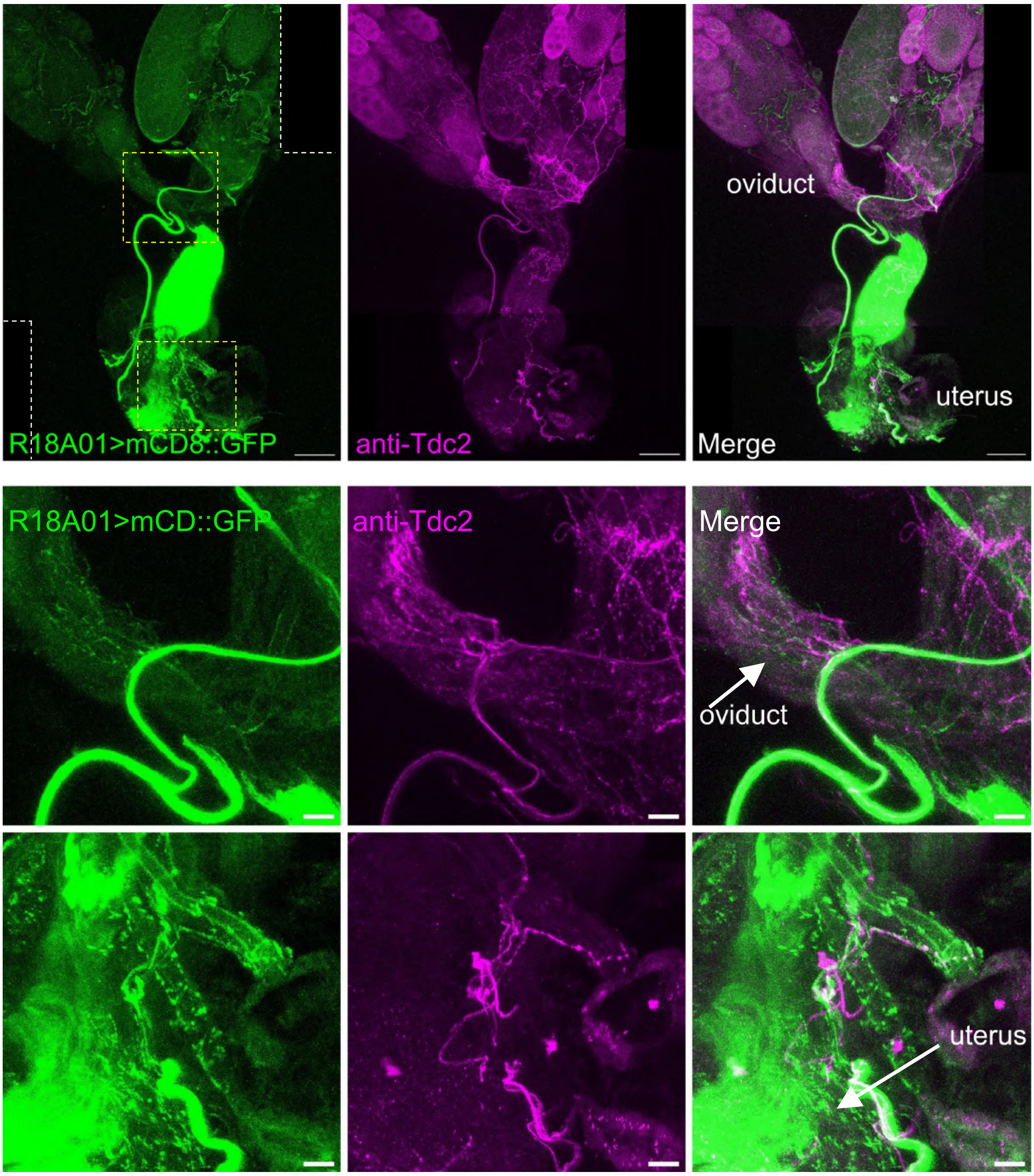


**Fig S13. Fig. S13. *R18A01>* is not strongly expressed in tyraminergic/octopaminergic neurons innervating the oviduct or uterus.** Top row, *R18A01>* is not strongly or unequivocally expressed in tyraminergic/octopaminergic neurons that innervate the ovary and oviduct. Depicted is a montage of two maximum intensity projections of confocal slices obtained from a dissected reproductive tract from a mated female of the genotype *2xR18A01>mCD8::GFP* and stained with with anti-GFP (green) and anti-Tdc2 (magenta), which marks tyraminergic/octopaminergic neurons. Dotted lines indicate regions not covered by the images. Same image as Fig. 6A, reproduced here for reference. Middle row, insert (top yellow dotted line in (B)), depicting the lateral oviducts and their junction with the common oviduct. No clear colocalization is observed. Bottom row, insert (bottom yellow dotted line in (B)), depicting the uterine region. Many neuromuscular junction-like termini are obsered in the uterine wall (region denoted by the arrow), which are negative for anti-Tdc2. Scale bars: (top panels) 100 µm; (bottom panels) 20 µm.

**Supplementary Figure S14.**

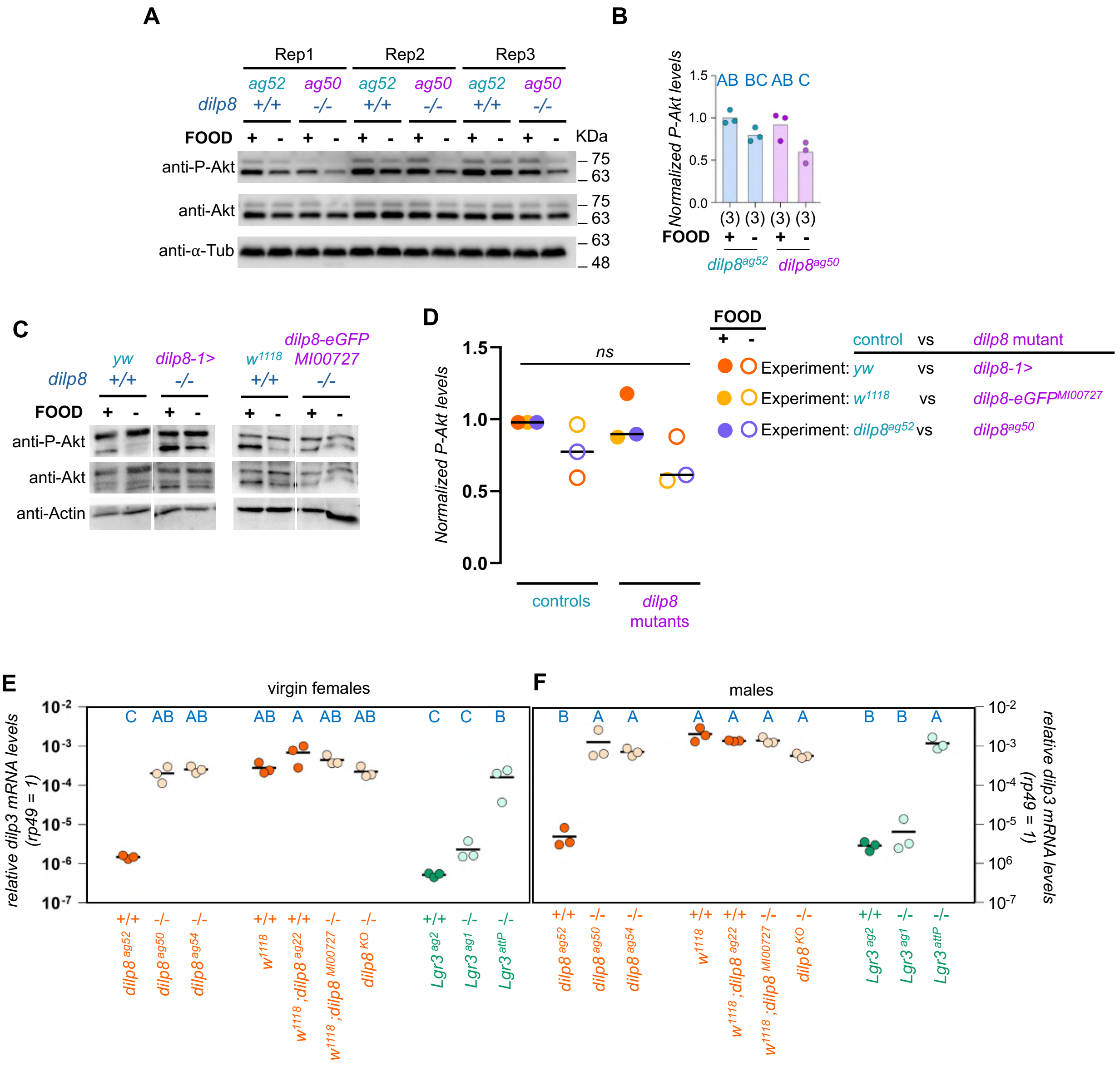


**Fig S14. Inconsistent effects of *dilp8* mutation on P-Akt levels. (A-D)** Here, we assayed for a systemic increase in insulin signaling in the body (head and thorax) of *dilp8* mutants relative to control animals using an antibody against phosphorylated Akt [pAkt (Ser473), equivalent to Ser405 in *Drosophila* (Yang et al. 2006)], which increases with insulin signaling (Burgering and Coffer 1995; Franke et al., 1995; Scanga et al., 2000; Yang et al. 2006). Control animals starved in 2% agar-water plates had a visible reduction in pAkt levels relative to well-fed animals (kept on yeast-food), as expected. Whereas yeast-fed *dilp8* mutants had similar levels of pAkt as controls, starved mutants had variable levels of pAkt relative to controls: *dilp8*$^{ag50}$ knockout mutants had statistically significantly reduced pAkt levels relative to fed controls and mutants, but not to starved *dilp8*$^{ag52}$ controls. These results were however not fully reproducible when two other *dilp8* mutants (*dilp8-eGFPMI00727* and *dilp8-1>*) were compared with their respective controls. One of them showed a similar effect upon starvation to *dilp8*$^{ag50}$, while the other mutant showed no effect upon starvation. The results are thus inconclusive, and suggest that if Dilp8 regulates systemic insulin signaling, this regulation is highly dependent on the genetic background. Alternatively, Dilp8 is not associated with systemic insulin signaling regulation and the variations in pAkt levels we detected are associated with other uncontrolled factors. **(A)** Western blots showing phosphorylation levels of Akt at serine 305 (P-Akt) are reduced upon 24 h starvation (in 2% agar-water plates) in the body (head and thorax) of control (*dilp8*$^{ag52}$)

and *dilp8* mutants (*dilp8*$^{ag50}$) virgins. Three independent repeats (Rep1-3) are shown. Food + and -, represent fed and starved conditions, respectively. Akt, total Akt. Alpha-Tubulin, loading control. **(B)** Quantification of both P-Akt bands relative to total Akt (both bands) depicted in (A). **(C)** Same as (A), but different control and *dilp8* mutant genotypes. The effect of starvation on some P-Akt bands is less prominent (left blots) or more pronounced (right blots), depending on the *dilp8* mutant flies analyzed. Actin, loading control. **(D)** Quantification of P-Akt relative to total Akt in three different experiments comparing different control and *dilp8* mutant genotypes. Each dot represents an experiment. The two experiments with a single repeat are depicted in (C). The average of these three repeats of the *dilp8*$^{ag52}$ vs *dilp8*$^{ag50}$ experiment depicted in (A) is used. Horizontal bar, average. All quantifications are normalized to their respective fed control. Fed animals, filled circles. Starved animals, empty circles. *ns,* $P > 0.05$, ANOVA and Tukey's HSD test. **(E-F)** Previous reports had shown that ectopic Dilp8 expression led to a reduction in the level of Dilp3 protein in adults of unspecified sex and mating status (Liao and Nässel 2020) and, complementary, that *dilp3* mRNA is slightly upregulated by a factor of 2.6 fold in 3-day old virgins in non-backcrossed *dilp8-eGFP*$^{MI00727}$ mutants relative to *w*$^{1118}$ animals (Li et al., 2023). To confirm these findings in a more controlled and homogeneous background, we performed qRT-PCR assays of whole virgin females from our previously published *dilp8* allele series (Heredia et al. 2021; Fernandez-Acosta et al., 2025). **(E)** Five-day old female virgins or **(F)** males homozygous for different *dilp8* or *Lgr3* mutant (-/-) alleles and their respective controls (+/+), show variable levels of *dilp3* mRNA, suggesting that they are responding to something else, possibly genetic background, and that it is not necessarily related to the role of Dilp8 and Lgr3 in ovulation control. Each dot represents a biological replicate. (B,E,F) Same blue letters, $P > 0.05$, ANOVA and Tukey's HSD test. Complete blots for (A,C) are shown in **Fig. S15**.

**Supplementary Figure S15.**

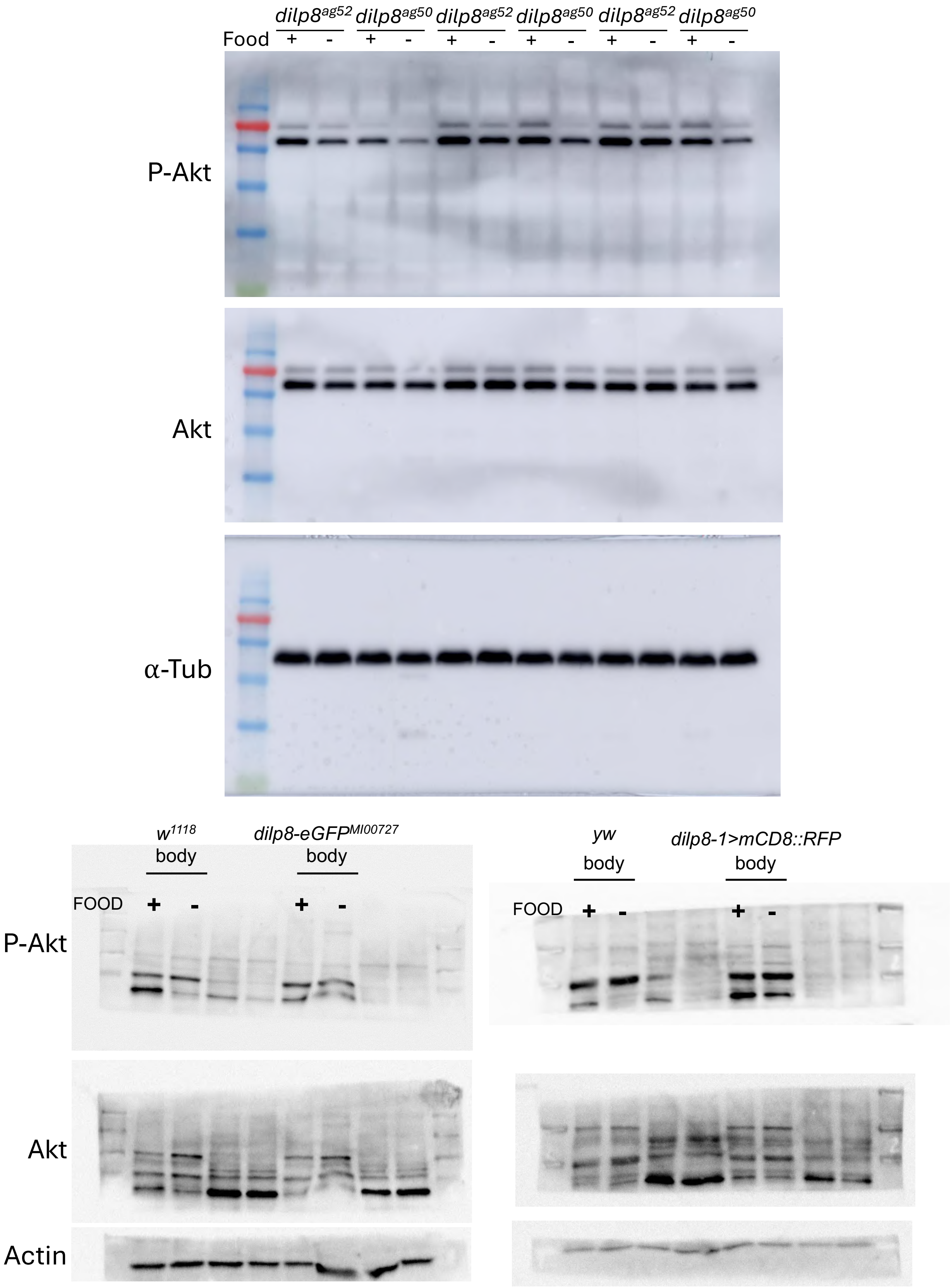


**Fig S15.** Full western blots showing anti-Akt phosphoylated at Ser505 (P-Akt), anti-total Akt (Akt), and anti-alpha-Tubulin (α-Tub) detection in the bodies (head and thorax) of fed (FOOD +) and starved (FOOD -) flies of the indicated genotypes. The red molecular marker migrates at 75 KDa.

**Supplementary Figure S16.**

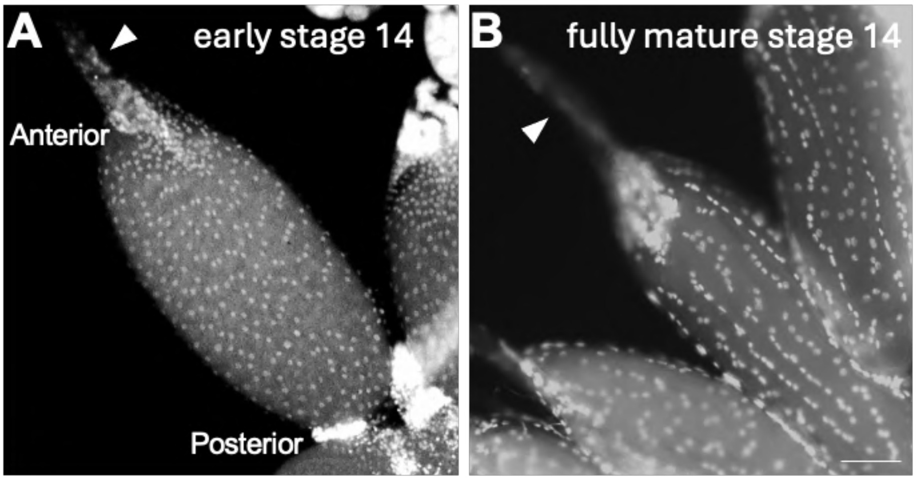


**Fig S16. Identification and classification of mature follicles in *D. melanogaster* ovaries. (A)** Ovaries were fixed and stained with DAPI to visualize nuclear organization. Mature follicles were identified by the absence of nurse cell nuclei in the anterior region and by their long dorsal appendage (arrowhead). Stage 14 follicles were classified as 'early' if their nuclei were randomly distributed and as **(B)** 'fully mature' if nuclei were aligned in a row. Scale bar: 50 μm.
